# Riboflavin metabolism shapes FSP1-driven ferroptosis resistance

**DOI:** 10.1101/2025.08.05.668651

**Authors:** Vera Skafar, Izadora de Souza, Ancely Ferreira dos Santos, Florencio Porto Freitas, Zhiyi Chen, Merce Donate, Palina Nepachalovich, Biplab Ghosh, Juliane Tschuck, Apoorva Mathur, Ariane Ferreira Nunes Alves, Jannik Buhr, Camilo Aponte-Santamaría, Werner Schmitz, Martin Eilers, Jessalyn Ubellacker, Ulrich Elling, Hellmut G Augustin, Kamyar Hadian, Svenja Meierjohann, Betina Proneth, Marcus Conrad, Maria Fedorova, Hamed Alborzinia, José Pedro Friedmann Angeli

## Abstract

Membrane protection against oxidative insults is achieved by the concerted action of glutathione peroxidase 4 (GPX4) and endogenous lipophilic antioxidants such as ubiquinone and vitamin E. Deficiencies in these protective systems lead to an increased propensity to phospholipid peroxidation and ferroptosis. More recently, ferroptosis suppressor protein 1 (FSP1) was identified as a critical ferroptosis inhibitor acting via regeneration of membrane-embedded antioxidants. Yet, regulators of FSP1 are largely uncharacterised, and their identification is essential for understanding the mechanisms buffering phospholipid peroxidation and ferroptosis. Here, we conducted a focused CRISPR-Cas9 screen to uncover factors influencing FSP1 function, identifying riboflavin (vitamin B₂) as a new modulator of ferroptosis sensitivity. We demonstrate that riboflavin, unlike other vitamins that act as radical-trapping antioxidants, supports FSP1 stability and the recycling of lipid-soluble antioxidants, thereby mitigating phospholipid peroxidation. Furthermore, we show that the riboflavin antimetabolite roseoflavin markedly impairs FSP1 function and sensitises cancer cells to ferroptosis. Thus, we uncover a direct and actionable role for riboflavin in maintaining membrane integrity by promoting membrane tolerance to lipid peroxidation. Our findings provide a rational strategy to modulate the FSP1-antioxidant recycling pathway and underscore the therapeutic potential of targeting riboflavin metabolism, with implications for understanding the interaction of nutrients and their contributions to a cell’s antioxidant capacity.

## Main text

Cellular membranes are crucial structural components, acting as dynamic barriers. Membrane lipid constituents are nevertheless vulnerable to oxidative damage, a process known as lipid peroxidation (LPO)^1^. LPO disrupts membrane integrity and executes ferroptosis, a type of regulated cell death implicated in a growing number of (patho)physiological processes, including cancer, neurodegeneration, and ischemia-reperfusion injury^2^.

Ferroptosis is predominantly inhibited by glutathione peroxidase 4 (GPX4)^3,4^, which reduces phospholipid hydroperoxides using glutathione. Additionally, radical-trapping antioxidants like ubiquinone (CoQ10), vitamin E, and vitamin K are crucial in suppressing propagation of lipid peroxidation^5^. More recently, ferroptosis suppressor protein 1 (FSP1) emerged as a key player in resisting ferroptosis^6,7^. Unlike GPX4, FSP1 regenerates quinone-like antioxidants such as ubiquinone and vitamin K^8^ using NAD(P)H as an electron donor, thereby halting the radical chain reaction required for ferroptosis execution. Although FSP1’s enzymatic mechanism is well understood, actionable factors regulating its function are largely uncharacterised. Identifying these regulators is vital for comprehending how cells maintain membrane redox homeostasis and withstand stress-induced ferroptosis, potentially revealing new therapeutic strategies for pathological conditions in which ferroptosis plays a significant role.

Here we used a CRISPR-Cas9 screen to identify regulators of FSP1 function, revealing riboflavin (vitamin B2) to be a modulator of ferroptosis sensitivity. Riboflavin, best known as a precursor for flavin adenine dinucleotide (FAD) and flavin mononucleotide (FMN), is essential for redox biology. We show that riboflavin directly supports the stability and activity of FSP1, enhancing its capacity to recycle lipid-soluble antioxidants and mitigate phospholipid peroxidation. Unlike other vitamins that act as direct radical-trapping antioxidants, riboflavin uniquely facilitates enzymatic recycling and is positioned upstream in the cascade, promoting ferroptosis resistance and preserving membrane integrity.

By revealing this novel role for riboflavin in regulating FSP1 and lipophilic antioxidant recycling, our findings expand our understanding of ferroptosis regulation and provide a framework for therapeutic strategies targeting the FSP1-antioxidant axis in cancer and other ferroptosis-associated diseases.

### Focused CRISPR-based screen uncovers new regulators of FSP1

We previously demonstrated that cells lacking GPX4 can survive and proliferate indefinitely when FSP1 activity is robust, either through naturally elevated FSP1 expression or enforced overexpression^6^ (Fig. 1a). Building on this observation, we generated FSP1-dependent HT1080 cells where GPX4 was deleted in an FSP1-overexpressing background (HT1080^GPX4KO/FSP1OE^) (Fig. 1b). HT1080^GPX4KO/FSP1OE^ cells readily undergo cell death upon FSP1 inhibition, which can be rescued by co-treatment with the ferroptosis inhibitor liproxstatin-1 (Lip1) (Fig. 1a, c).

**Figure 1.**
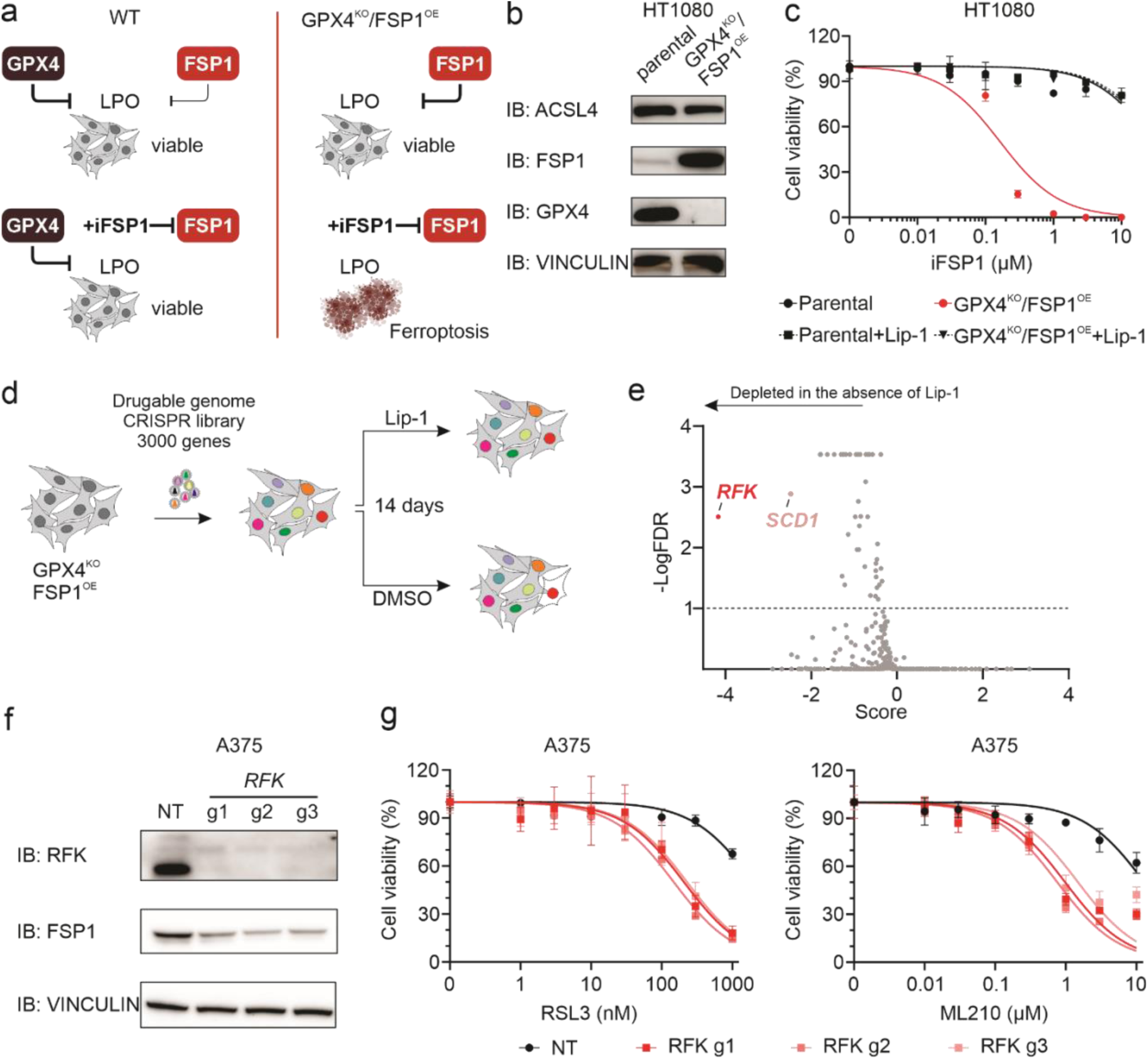
Identification of factors supporting FSP1 function. **a,** Schematic representation of the FSP1-dependent model used to identify factors supporting FSP1 function. Upper left: the primary protective system against lipid peroxidation (LPO) is the enzyme GPX4, complemented by FSP1. Lower left: cells treated with an FSP1 inhibitor (iFSP1) can rely on GPX4 activity to survive. Upper right: FSP1-dependent HT1080 cells (HT1080^GPX4KO/FSP1OE^). In HT1080 cells, as in many others, knocking out GPX4 induces ferroptosis due to insufficient endogenous FSP1 levels to compensate for GPX4 loss. However, cell survival can be rescued by overexpressing FSP1. Lower right: Upon pharmacological inhibition of FSP1 (iFSP1 treatment), cells undergo ferroptosis as they solely rely on FSP1 function for survival. **b**, Immunoblot (IB) analysis of ACSL4, FSP1, GPX4, and vinculin in HT1080 parental and HT1080^GPX4KO/FSP1OE^ cells. **c**, Dose-dependent toxicity of an FSP1 inhibitor (iFSP1) in HT1080 parental and HT1080^GPX4KO/FSP1OE^ cell lines. Cell viability was monitored using Alamar blue after 24 h of treatment. Where indicated, cells were treated with the ferroptosis inhibitor Lip-1 (500 nM). **d**, Schematic representation of the screening strategy used to identify novel factors supporting FSP1 function in the previously-described FSP1-dependent cellular model (see Fig. 1a). HT1080^GPX4KO/FSP1OE^ cells were transduced with a gRNA library targeting approximately 3000 genes and selected over 7 days in the presence of the ferroptosis inhibitor Lip-1 (500 nM). Subsequently, cells proliferate with or without Lip-1 supplementation for additional 14 days. **e,** Graphical representation of the results from two independently performed CRISPR screens; plot depicts the score calculated using the MaGeCK package (x-axis) and the –Log false discovery rate (y-axis). Riboflavin kinase (RFK) and stearoyl-CoA desaturase-1 (SCD1) were identified as potentially robust candidates to modulate FSP1 function. **f**, Immunoblot (IB) analysis of RFK, FSP1, and vinculin in A375 cells transduced with either a non-targeting control (NT) or three different RFK-targeting sgRNAs. **g**, Dose-dependent toxicity of RSL3 and ML210 in A375 cells transduced with either a non-targeting control (NT) or three different RFK-targeting sgRNAs. Cell viability was monitored after 72 h of treatment.

We reasoned that this cellular model could be combined with CRISPR-based genetic screening to identify factors contributing to FSP1 function. Therefore, we performed a focused CRISPR screen (targeting approximately 3000 potential druggable genes) in this line, comparing conditions with and without Lip1. We posited that genetic perturbations impairing FSP1 expression or activity would induce ferroptosis, which Lip1 should rescue (Fig. 1d)^6^. Through this approach, we identified several genes whose loss significantly impacted FSP1-dependent ferroptosis resistance, with the two top hits being stearoyl-CoA desaturase (SCD1) and riboflavin kinase (RFK) (Fig. 1e and Extended Table 1). Loss of SCD1 is known to increase ferroptosis sensitivity by raising the PUFA/MUFA ratio^9^. Consistent with this, pharmacological inhibition of SCD1 readily triggered ferroptosis in HT1080^GPX4KO/FSP1OE^ cells (Extended Data Fig. 1a, b), likely due to an elevated pool of oxidisable substrates overwhelming FSP1’s protective capacity. These findings suggest that cells with a high PUFA/MUFA ratio derive limited benefit from FSP1 activity. However, SCD1 depletion did not appear to directly impair FSP1 function.

Given these results, we focused on RFK and its role in regulating FSP1. RFK is a key enzyme, phosphorylating riboflavin to generate flavin mononucleotide (FMN), a central step in the production of flavin adenine dinucleotide (FAD)^10^—a co-factor essential for the activity of flavoproteins, including FSP1^11^. While FSP1’s dependence on FAD implies a direct link to RFK, the extent to which RFK deficiency affects FSP1 levels and ferroptosis resistance is unknown. Using A375 cells, in which FSP1 confers strong protection against GPX4 inhibitors (GPX4i) such as RSL3 and ML210, we found that CRISPR-mediated deletion of RFK leads to a robust loss of RFK expression and a decrease in FSP1 protein levels (Fig. 1f). This reduction renders RFK-deficient cells highly sensitive to GPX4i, highlighting a previously uncharacterised dependency of ferroptosis resistance on RFK (Fig. 1g). We obtained similar results using HT1080^GPX4KO/FSP1OE^ cells, where we find that the absence of RFK impairs viability and induces ferroptosis (Extend Data 1 c-f). Our findings establish RFK as an actionable upstream regulator of FSP1 functionality and ferroptosis resistance.

### FAD deficiency disrupts FSP1 function and promotes ferroptosis susceptibility

Interestingly, we observed that increased sensitivity to GPX4 inhibitors (GPX4i) in cells transduced with sgRNA targeting RFK was lost over time. FAD biosynthesis is a multi-step process beginning with riboflavin uptake via members of the SLC52A family of solute carriers (SLC52A1, SLC52A2, and SLC52A3). Once inside the cell, riboflavin is phosphorylated by RFK to generate FMN, which is subsequently adenylated by flavin adenine dinucleotide synthetase 1 (FLAD1) to produce FAD (Fig. 2a). Consistently, analysis of the DepMap database (www.depmap.org) reveals that RFK loss is poorly tolerated by most cell types (Fig. 2b). This intolerance was evident in our cultures, where unedited cells rapidly outcompeted RFK-deficient cells (Extended Data, Fig. 2a). These challenges impeded further experiments and prompted us to explore whether other enzymes generating the flavin co-factors could be targeted.

**Figure 2.**
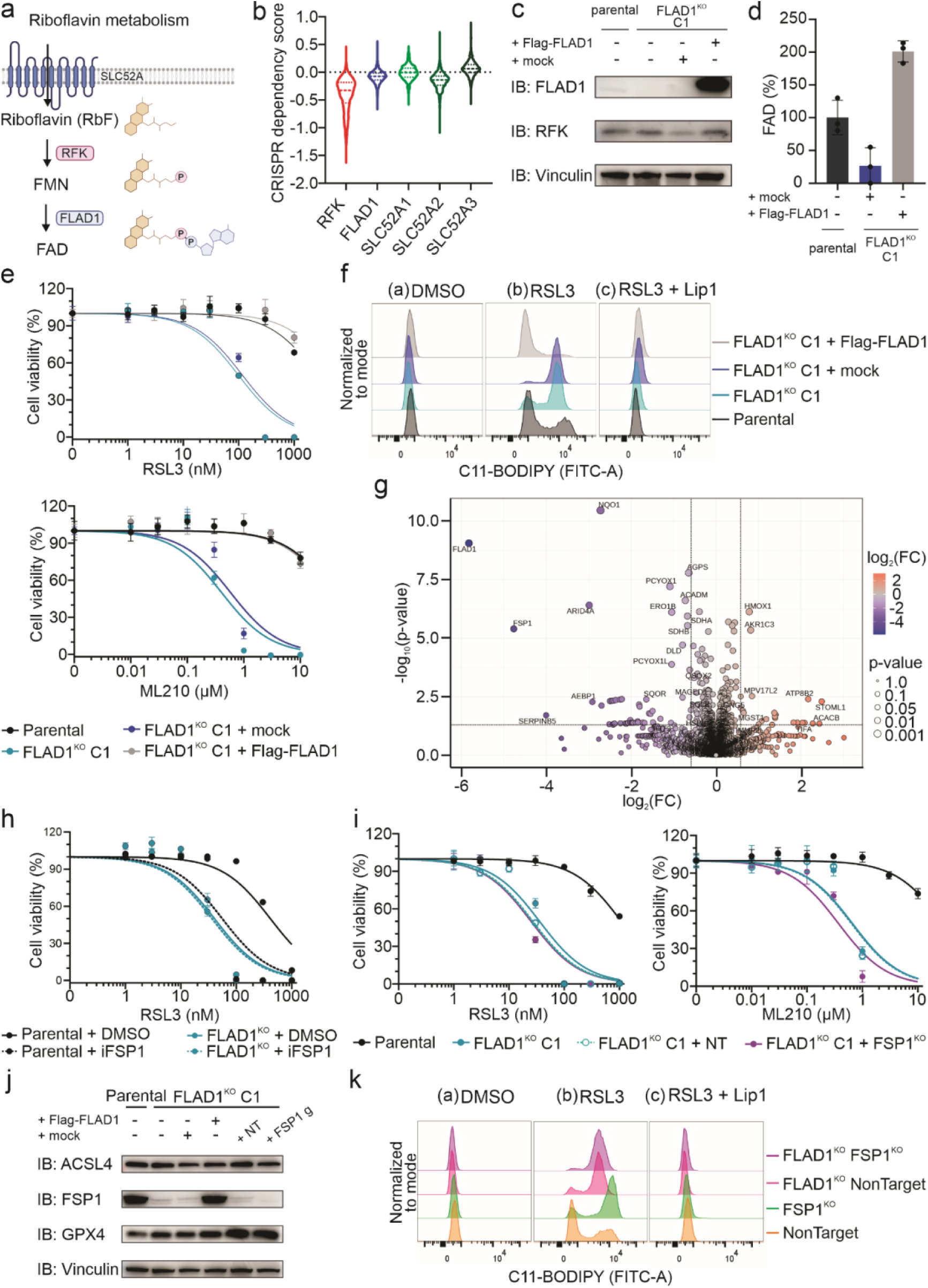
FAD deficiency disrupts FSP1 function and promotes ferroptosis susceptibility. **a**, Schematic representation of flavin mononucleotide (FMN) and flavin adenine dinucleotide (FAD) biosynthesis from riboflavin (RbF). Riboflavin is first phosphorylated to form FMN in a reaction catalyzed by riboflavin kinase (RFK, encoded by *RFK*). FMN is then adenylated to form FAD in a reaction catalyzed by FAD synthase (FADS, encoded by *FLAD1*). **b**, CRISPR dependency scores of *RFK*, *FLAD1,* and *SLC52A1-3* perturbations across a panel of human cancer cell lines (https://depmap.org/portal/, version 23Q2). **c**, Immunoblot (IB) analysis of FLAD1, RFK, and vinculin in A375 parental, A375 FLAD1^KO^ single clone 1 (C1) and A375 FLAD1^KO^ C1 cells stably overexpressing either an empty vector (mock) or Flag-FLAD1 (addback). **d**, Relative quantification of FAD in A375 parental and A375 FLAD1^KO^ C1 cells stably overexpressing either an empty vector (mock) or Flag-FLAD1. **e**, Dose-dependent toxicity of RSL3 and ML210 in A375 parental, A375 FLAD1^KO^ single clone 1 (C1) and A375 FLAD1^KO^ C1 cells stably overexpressing either an empty vector (mock) or Flag-FLAD1. Cell viability was monitored using Alamar blue after 72 h of treatment. **f**, Lipid peroxidation evaluated by C11-BODIPY 581/591 staining of A375 parental, A375 FLAD1^KO^ single clone 1 (C1) and A375 FLAD1^KO^ C1 cells stably overexpressing either an empty vector (mock) or Flag-FLAD1. Cells were treated with DMSO, RSL3 (200 nM), or RSL3 (200 nM) + Lip-1 (500 nM) for 6 h. **g**, Volcano plot of differentially-expressed proteins between A375 FLAD1^KO^ C1 and A375 FLAD1^KO^ C1 cells stably overexpressing Flag-FLAD1. Quantified proteins are plotted based on their fold change (FC: FLAD1^KO^ C1/FLAD1^KO^ C1 Flag-FLAD1^OE^). **h**, Dose-dependent toxicity of RSL3 in A375 parental and A375 FLAD1^KO^ cells in the absence or presence of an FSP1 inhibitor (iFSP1 2 µM). Cell viability was monitored using after 72 h of treatment. **i**, Dose-dependent toxicity of RSL3 and ML210 in A375 parental, A375 FLAD1^KO^ single clone 1 (C1) and A375 FLAD1^KO^ C1 transduced with either a non-targeting control (NT) or a FSP1-targeting sgRNAs (FSP1^KO^). Cell viability was monitored after 72 h of treatment. **j**, Immunoblot (IB) analysis of ACSL4, FSP1, GPX4, and vinculin in A375 parental, A375 FLAD1^KO^ C1, A375 FLAD1^KO^ C1 cells stably overexpressing an empty vector (mock) or Flag-FLAD1 and A375 FLAD1^KO^ C1 transduced with either a non-targeting control (NT) or a FSP1-targeting sgRNAs (FSP1^KO^). **k**, Lipid peroxidation evaluated by C11-BODIPY 581/591 staining of A375 parental, A375 FLAD1^KO^ single clone 1 (C1), and A375 FLAD1^KO^ C1 cells transduced with either a non-targeting control (NT) or a FSP1-targeting sgRNAs (FSP1^KO^). Cells were treated with DMSO, RSL3 (200 nM), or RSL3 (200 nM) + Lip-1 (500 nM) for 6 h.

Notably, unlike RFK, loss of FLAD1—the enzyme responsible for the final step in FAD biosynthesis—appears to be better tolerated by cells, likely because essential FMN-dependent proteins remain functional. Therefore, we knocked out FLAD1 in A375 cells and HT1080^GPX4KO/FSP1OE^ cells and single clones thereof (Fig. 2c and Extended Data, Fig. 2b-g), which allowed us to establish an isogenic pair of FLAD1-proficient and -deficient cell lines (Fig 2c,d). Using this model, we find that loss of FLAD1 leads to markedly increased sensitivity to GPX4 inhibition (Fig. 2e) as well as to lipid peroxidation (Fig. 2f). We saw similar effects in the HT1080^GPX4KO/FSP1OE^ model (Extended Data, Fig. 2h, j).

Loss of flavin cofactors impairs the stability of the flavoproteome and exerts a broad metabolic effect, including impacting lipid metabolism^12^. Therefore, we aimed to determine if the effect— loss of redox homeostasis—is FSP1-specific or more general. A whole proteomic analysis of the FLAD1 isogenic pair confirmed a reduced abundance of flavoproteins. Yet interestingly, FSP1 and NQO1 are the most strongly depleted flavoproteins in FLAD1-deficient cells (Fig. 2g), an effect independent of mRNA abundance (Extended Data, Fig. 2k). Building on this observation, we validated the effect in other cell lines, showing that genetic loss of FLAD1 leads to a substantial loss of FSP1 and is accompanied by increased sensitivity to ferroptosis Extended Data Fig. 2l, m).

Molecular dynamic simulations of FSP1 with and without FAD were carried where the RMSD analysis shows that the absence of FAD increases protein backbone instability (Extended Data, Fig. 2n). RMSF profiles reveal more significant residue fluctuations, especially in residues 282-300, which interact with FAD (Extended Data, Fig. 2o) following our previous report ^13^. Overall, these findings establish that FAD is essential not only for activity but also for FSP1 stability. Building on this observation, we validated the effect in other cell lines, showing that genetic loss of FLAD1 leads to a substantial loss of FSP1 and is accompanied by increased sensitivity to ferroptosis Extended Data Fig. 2l, m). Using genetic and pharmacological approaches, we find that loss or inhibition of FSP1 in FLAD1-deficient cells causes no further sensitisation to lipid peroxidation and ferroptosis induced by GPX4i (Fig 2h-k and Extended Data Fig. 2p). Lastly, loss of FLAD1 appears to sensitise cells to GPX4i specifically (Fig 2h-k): other ferroptosis inducers, such as Erastin and BSO, or other tested cytotoxic agents encompassing a diverse variety of modes of action (Extended Data Fig. 2p, q) exert no enhanced effects in the absence of FLAD1. These findings establish FLAD1 as a key regulator of FSP1 activity and ferroptosis sensitivity.

### Riboflavin availability as a central determinant of ferroptosis resistance

Having established a functional link between intracellular flavin metabolism and ferroptosis resistance via FSP1, we hypothesised that the availability of riboflavin—the main precursor for FAD—can influence ferroptosis sensitivity. To test this, we examined proteomic changes in cells cultured under riboflavin-deficient conditions (Fig. 3a and Extended Fig. 3a-c). As in FLAD1-deficient cells, we observe an overall loss of flavoproteins. However, the changes appear more extensive, presumably because FMN-containing proteins are also affected Extended Fig. 3d-e). Notably, FSP1 emerged again as the most downregulated protein in riboflavin-deficient conditions (Fig. 3a, b), underscoring the critical role of FAD in FSP1 stability. Riboflavin withdrawal is also accompanied by a characteristic upregulation of NRF2 target genes (e.g., AKR1Cs, TXNRD1, GCLM).

**Figure 3.**
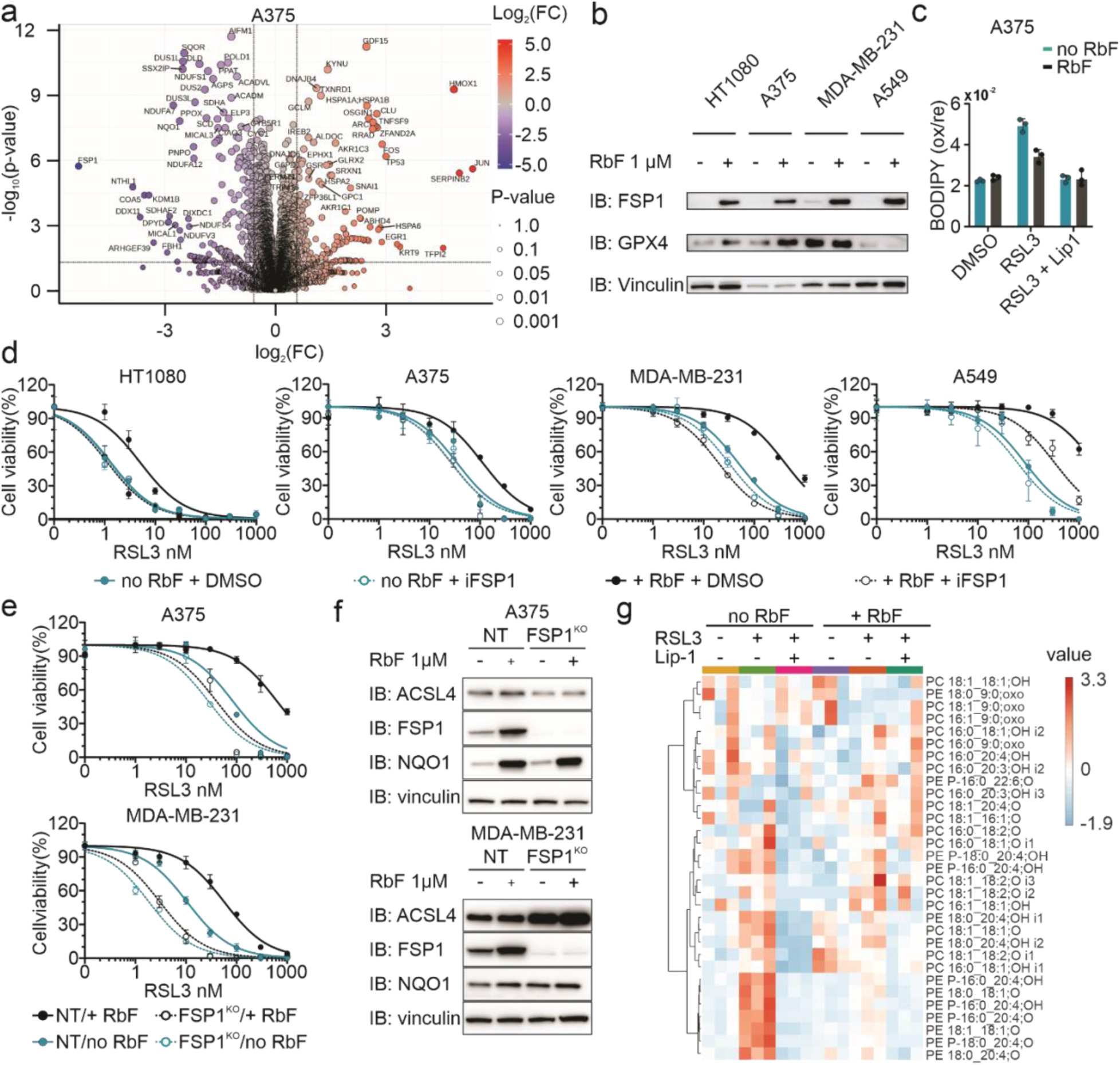
Riboflavin availability as a central determinant of ferroptosis resistance. **a**, Volcano plot of quantified proteins showing their change in A375 parental cells cultured in riboflavin-deficient medium for 96 h. Proteins are plotted based on their fold change (FC: riboflavin deficient/normal). **b**, Immunoblot (IB) analysis of FSP1, GPX4, and vinculin in HT1080, A375, MDA-MB-231, and A549 parental cell lines after 96 h of growth in riboflavin-deficient medium or medium supplemented with 1 µM riboflavin. **c**, Lipid peroxidation evaluated by C11-BODIPY 581/591 staining of A375 parental cell line cultured for 72 h in riboflavin-deficient medium or medium supplemented with 1 µM riboflavin and after treatment with DMSO, RSL3 (200 nM), or RSL3 (200 nM) + Lip-1 (500 nM) for 6 h. **d**, Dose-dependent toxicity of RSL3 in the absence or presence of an FSP1 inhibitor (iFSP1 3 µM) in HT1080, A375, MDA-MB-231, and A549 parental cell lines pre-cultured in riboflavin-deficient medium or medium supplemented with 1 µM riboflavin for 48 h. Cell viability was monitored using Alamar blue after 96 h of treatment. **e**, Dose-dependent toxicity of RSL3 in the absence or presence of an FSP1 inhibitor (iFSP1 3 µM) in A375 and MDA-MB-231 cells transduced with either a non-targeting control (NT) or an FSP1-targeting sgRNAs pre-cultured in riboflavin-deficient medium or medium supplemented with 1 µM riboflavin for 48 h. Cell viability was monitored after 96 h of treatment. **f**, Immunoblot (IB) analysis of ACSL4, FSP1, NQO1, and vinculin in A375 and MDA-MB-231 cells transduced with either a non-targeting control (NT) or an FSP1-targeting sgRNAs cultured in riboflavin-deficient medium or medium supplemented with 1 µM riboflavin for 96 h. **g**, Epilipidomics analysis of A375 parental cells pre-cultured in riboflavin-deficient medium or medium supplemented with 1 µM riboflavin for 72 h after treatment with DMSO, RSL3 (200 nM), or RSL3 (200 nM) + Lip-1 (500 nM) for 6 h.

Culturing cells without riboflavin for 72 hours markedly increases their susceptibility to lipid peroxidation (Fig. 3c and Extended Fig. 3f) and enhances cell death upon GPX4 inhibition (GPX4i) (Fig. 3d and Extended Fig. 3g). We could rule out the sensitisation to ferroptosis upon riboflavin withdrawal by a direct antioxidant role of riboflavin (Extended Data Fig. 3h). These effects are not limited to A375 cells: three additional cancer cell lines from different tissue origins exhibit similar responses (Fig. 3b, d). Consistent with an FSP1-dependent mechanism, combining GPX4i with an FSP1 inhibitor strongly sensitised cells under riboflavin-replete conditions but had negligible effects in the absence of riboflavin (Fig. 3d). We further corroborated these effects in FSP1-deficient A375 and MDA-MB-231 cell lines. Upon GPX4 inhibition, only cells cultured with riboflavin displayed further ferroptosis sensitisation following FSP1 loss (Fig. 3e, f). These results were recapitulated in HT1080^GPX4KO/FSP1OE^ and A375 cells where genetic deletion of the main riboflavin transporter, namely SLC52A2, disrupted FSP1 function and sensitised cells to GPX4i (Extended Data Fig. 4a-l). These findings indicate that, under riboflavin-poor conditions, cells become ferroptosis-sensitive primarily through an FSP1-dependent pathway.

To directly demonstrate that riboflavin supports protection against LPO, we analysed the epilipidome of cells treated with GPX4i under riboflavin-replete and riboflavin-deficient conditions (Fig. 3g). Our detailed analysis revealed a rapid and specific accumulation of oxidised phosphatidylethanolamine (PE) species^14^ upon GPX4 inhibition in riboflavin-starved cells. Critically, treatment with Lip-1 fully reverses the formation of oxidised lipids. Together, these results establish a direct link between riboflavin and membrane antioxidant capacity.

Given our results under riboflavin-depleted conditions, we asked whether moderate variations in riboflavin concentration would similarly affect sensitivity to GPX4i. Notably, human plasma riboflavin concentrations typically range from 10 to 20 nM^15^, while standard cell culture media like RPMI and DMEM contain around 500 and 1000 nM, respectively. More advanced media, such as Plasmax^16^ and HPLM^17^, also include significantly higher riboflavin levels at 300 and 500 nM, respectively. We found that physiological riboflavin levels (≤20 nM) markedly reduce FSP1 expression, whereas concentrations above 100 nM are sufficient to stabilise FSP1 and confer ferroptosis resistance (Extended Data, Fig. 4j). Altogether, our studies demonstrate that riboflavin availability is a central determinant of membrane repair capacity and determines FSP1 antioxidant recycling capacity.

**Figure 4.**
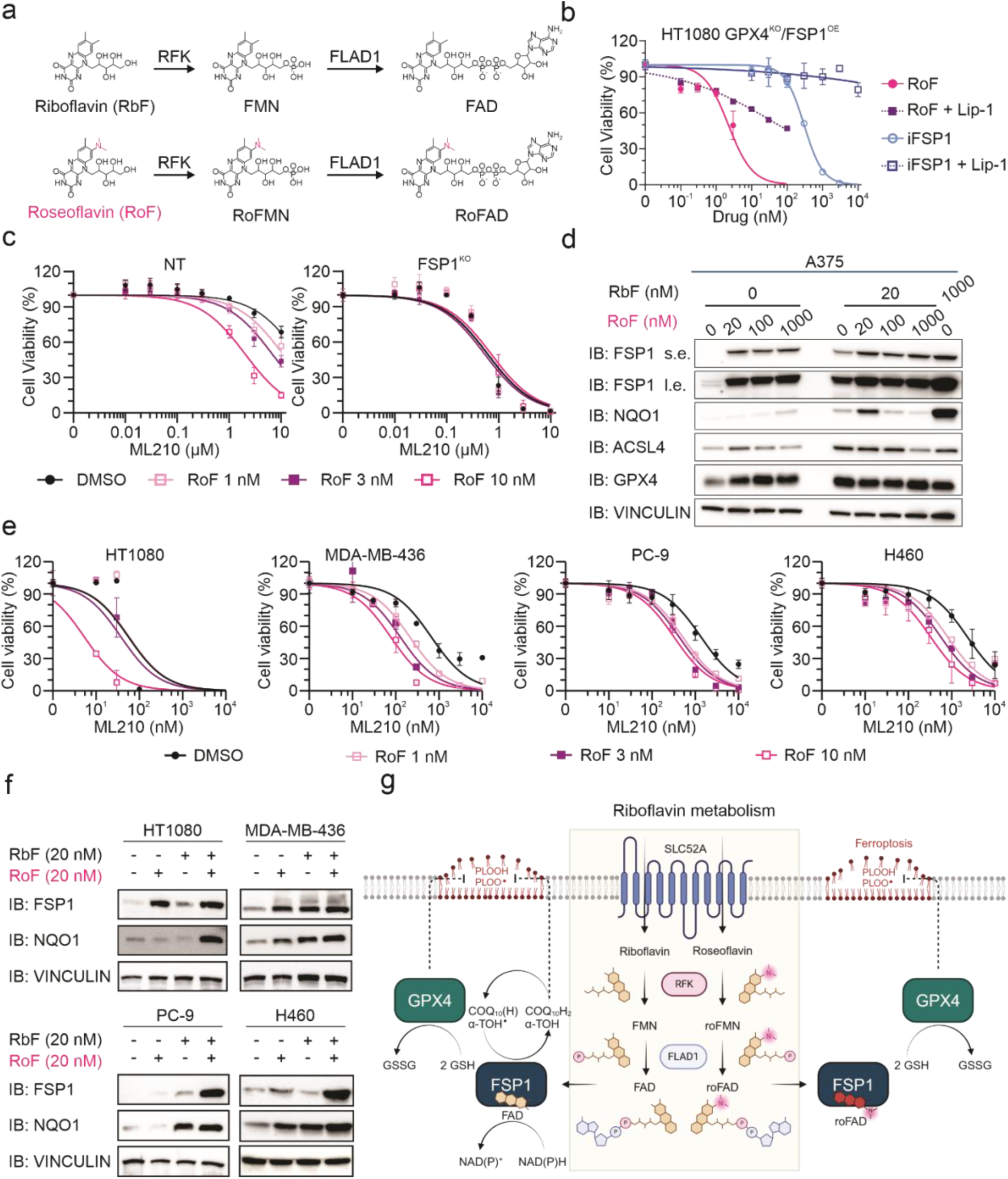
The riboflavin analogue roseoflavin disrupts FSP1 activity and promotes ferroptosis. **a**, Schematic representation of the metabolism of riboflavin (RbF) and its analog roseoflavin (RoF) by riboflavin kinase (encoded by *RFK*) and FAD synthase (encoded by *FLAD1*). **b**, Dose-dependent toxicity of roseoflavin (RoF) and iFSP1 in the absence or presence of Lip-1 (500 nM) in HT1080^GPX4KO/FSP1OE^. Cell viability was monitored using Alamar blue after 48 h of treatment. Cells were cultured in low-riboflavin medium (20 nM). **c**, Dose-dependent toxicity of the GPX4 inhibitor ML210 in A375 cells transduced with either a non-targeting control (NT) or a FSP1-targeting sgRNAs (FSP1^KO^) that were pre-treated with roseoflavin (RoF, 1, 3, and 10 nM) for 48 h. Cell viability was monitored after 48 h of ML210 treatment. Cells were cultured in low-riboflavin medium (20 nM). **d**, Immunoblot analysis (IB) of FSP1, NQO1, ACSL4, GPX4, and vinculin treated with roseoflavin (RoF 0, 20, 100, and 1000 nM) for 96 h in the absence or presence of riboflavin (RbF 20 nM). **e**, Dose-dependent toxicity of the GPX4 inhibitor ML210 in HT1080, MDA-MB-436, PC-9, and H460 parental cell lines pre-treated with roseoflavin (RoF, 1, 3, and 10 nM) for 48 h. Cell viability was monitored after 48 h of ML210 treatment. **f**, Immunoblot (IB) analysis of FSP1, NQO1, and vinculin in HT1080, MDA-MB-436, PC9, and H460 parental cell lines treated with roseoflavin (RoF, 20 nM) for 48 h in the absence or presence of riboflavin (RbF 20 nM). **g**, Riboflavin metabolism supports FSP1 function and promotes ferroptosis resistance. This is therefore a process that riboflavin analogs, such as roseoflavin can modulate.

### Riboflavin analogues disrupt FSP1 activity and promote ferroptosis

Our results suggest that pharmacologically targeting riboflavin metabolism, to disrupt its FSP1-protective branch, would sensitise cancer cells to ferroptosis. No selective inhibitors for any of the proteins involved in riboflavin uptake (SLC52A2) or its action toward FMN (RFK) and FAD (FLAD1) have been described. Nonetheless, bacteria from the genus *Streptomyces (e.g*. *davaonensis* and *cinnabarinus*) produce the riboflavin antimetabolite roseoflavin that exerts antimicrobial activity by binding to and disrupting riboflavin riboswitches. Roseoflavin has shown promise as a broad-spectrum antibiotic, but few studies have explored it as an anticancer agent. In eukaryotic cells, reseoflavin is thought to follow a similar metabolic route as riboflavin—being transported, phosphorylated, and adenylated (Fig. 4a) ^18^. The dimethylamino group on C8 of the isoalloxazine ring leads to the formation of an altered flavin cofactor that is believed to disrupt normal flavin-mediated electron transfer reactions.

Using our HT1080^GPX4KO/FSP1OE^ cells, we investigated whether roseoflavin influences FSP1-mediated ferroptosis protection. Notably, at physiologically relevant levels of riboflavin, roseoflavin induced ferroptosis within the single-digit nanomolar range (Fig. 4b). Moreover, roseoflavin affects the response to GPX4i only in FSP1-expressing cells (Fig. 4c and Extended Data Fig. 5 a, b); no sensitisation was detected in FSP1-deficient cells (Fig. 4c and Extended Data Fig. 5 c), indicating that the effect of roseoflavin is specific. Additionally, roseoflavin can restore FSP1 levels when cells are treated under low and physiological riboflavin conditions (Fig 4e), suggesting that the effect is likely on-target. Finally, the effect of roseoflavin was broadly reproduced in a larger panel of cell lines (Fig 4f), where treatment can restore FSP1 levels (Fig 4e). These actions reflect the ability of roseoflavin adenine dinucleotide (roFAD), like FAD, to stabilise FSP1; however, the modification of the isoalloxazine group destroys its oxidoreductase function.

Altogether, these results demonstrate that FSP1 activity can be effectively inhibited by riboflavin antimetabolites, offering additional opportunities to target ferroptosis protective mechanisms in cancer cells.

## Discussion

Here we reveal a critical role for riboflavin in governing ferroptosis via FSP1. Previous work established FSP1 as an essential safeguard against lipid peroxidation, complementing the activity of GPX4; however, the upstream factors regulating FSP1 functionality were largely unknown. By systematically dissecting FAD biosynthesis, we demonstrate that multiple nodes in the riboflavin– FMN–FAD pathway directly influence FSP1’s stability and enzymatic capacity to recycle lipophilic antioxidants. In particular, we highlight FLAD1 as a key enzyme in the terminal step of FAD production and reveal that its depletion specifically compromises FSP1, with broad implications for ferroptosis sensitivity and cell viability. These findings enhance our understanding of how cells depend on riboflavin metabolism to preserve membrane integrity under oxidative stress.

Crucially, we observed that standard tissue culture media supply riboflavin at concentrations far exceeding physiological plasma levels, potentially obscuring the impact of riboflavin availability on ferroptosis. Culturing cells at physiological or deficient riboflavin levels revealed pronounced changes in FSP1 expression and ferroptosis outcomes, underscoring how even moderate fluctuations in riboflavin can alter redox balance. This insight raises important considerations for mechanistic studies and potential clinical contexts in which dietary riboflavin or altered metabolism may drive vulnerability to ferroptosis-related diseases^19^.

Interestingly, a parallel can be drawn between the regulation of FSP1 by riboflavin and the regulation of GPX4 by selenium. It is well-established that selenium availability dictates GPX4 protein translation, with selenium metabolism pathways attracting significant interest as potential modulators of ferroptosis^20–22^. Similarly, our findings suggest that riboflavin—by serving as a precursor for FAD—determines the abundance and activity of FSP1 independently of mRNA levels. Importantly, both GPX4 and FSP1 highlight how two crucial regulators of ferroptosis cannot be understood solely by analysing the abundance of their mRNA. Instead, the availability of specific micronutrients—selenium for GPX4 and riboflavin for FSP1—is required for their proper translation and functionality. This establishes a broader principle where micronutrients and their metabolic pathways modulate protein functionality and ferroptosis susceptibility, suggesting significant new opportunities for therapeutic intervention.

Beyond elucidating a role for endogenous riboflavin metabolism, we highlight the value of riboflavin antimetabolites, such as roseoflavin, as potent modulators of FSP1 function. Owing to its structural similarity to riboflavin, roseoflavin likely shares efficient tissue distribution and demonstrates activity in the nanomolar range in physiologically relevant riboflavin concentrations—substantially lower than current FSP1 inhibitors. An additional advantage of roseoflavin lies in its uptake and metabolism, which depend on SLC52A2, RFK, and FLAD1. Mutations in these proteins that could affect resistance would also compromise riboflavin uptake and metabolism, ultimately preventing the production of essential cofactors like FAD and FMN, as recently exemplified in *Plasmodium falciparum*^23^. This dual impact makes it unlikely for cells to selectively develop resistance to roseoflavin without severely impairing riboflavin metabolisation and FAD production. In contrast, specific inhibitors of FSP1 are more prone to resistance mechanisms, such as mutations in FSP1 itself or activation of compensatory pathways, as these do not disrupt the broader metabolic network. Roseoflavin’s reliance on essential and conserved metabolic pathways could thus provide a significant therapeutic advantage.

Altogether, our results show that intracellular flavin metabolism is pivotal to FSP1’s protective function against phospholipid peroxidation, operating independently of GPX4. This framework constitutes a previously underappreciated approach for enhancing ferroptosis in cancer cells and other contexts where FSP1 supports survival. Moreover, our work unveils riboflavin’s role in the FSP1-driven recycling of lipophilic antioxidants, offering fundamental insight into the complex interaction of nutrients with important implications for understanding inconsistent outcomes in preclinical and clinical studies of antioxidant therapies ^24–26^.

## Acknowledgements

J.P.F.A. acknowledges the support of the Junior Group Leader program of the Rudolf Virchow Center, University of Würzburg and additional support from the Deutsche Forschungsgemeinschaft (DFG), FR 3746/3-1, FR 3746/5-1, FR 3746/6-1, CRC205 (INST 269/886-1), and TRR 387/1 (514894665); the EU-H2020 (ERC-Consolidator, DeciFERR); the Deutsche Jose Carreras Leukämie Stiftung (DJCLS 01 R/2022); and The São Paulo Research Foundation (2023/04397-4). Work in the Fedorova lab is supported by ‘‘Sonderzuweisung zur Unterstützung profilbestimmender Struktureinheiten’’ by the SMWK to TUD, TG70 by Sächsische Aufbaubank and SMWK, the measure is co-financed with tax funds based on the budget passed by the Saxon state parliament (to M.F.), Deutsche Forschungsgemeinschaft (FE 1236/5-1, FE 1236/8-1 to M.F.), and Bundesministerium für Bildung und Forschung (031L0315A, DEEP_HCC and 01EJ2205A, FERROPath to M.F.). We also want to acknowledge the German Cancer Research Center (DKFZ) Proteomics Core Facility. M.C. acknowledges support from DFG (CO 291/7-1, the Priority Program SPP 2306 [CO 291/9-1, #461385412; CO 291/10-1, #461507177], and the CRC TRR 353 [CO 291/11-1; #471011418]); the European Research Council (ERC) under the European Union’s Horizon 2020 research and innovation programme (grant agreement no. GA 884754) and the German Federal Ministry of Education and Research (BMBF) FERROPATH (01EJ2205B) to M.C. and B.P. Illustrations in Fig. 2a and 4g were created in BioRender. Skafar, V. (2025) https://BioRender.com/j20e353).

## Author Contributions

V.S., and I.S., performed most experiments with support from A.F.S., F.P.F. and M.D. Z.C. performed and analysed the CRISPR screen. P.N. and M.F performed epilipidomics analysis.. W.S. performed the analysis and interpretation of flavin quantification using HPLC-MS. B. G. contributed with the statistical analysis of proteomics data. S.M. designed and assisted with RT-qPCR analysis. A.M, A.F.N.A., J. B., C.A-S., carried molecular dynamic simulations to determine the impact of FAD loss in FSP1 stability. H.G.A., S.M., U. E., M.E., J. T., K. H., A. F. N. A., J. B., C.A., R.G.B., B.P., M.C., M.F., and H. A. contributed with reagent, critical information and/or platforms. J.P.F.A. initiated and supervised the project. All authors contributed with discussion and data interpretation and read and agreed on the paper’s content.

## Conflict of Interest

M.C. and B.P. are co-founders and shareholders of ROSCUE Therapeutics GmbH.

## Extended Figures

**Extended Figure 1.**
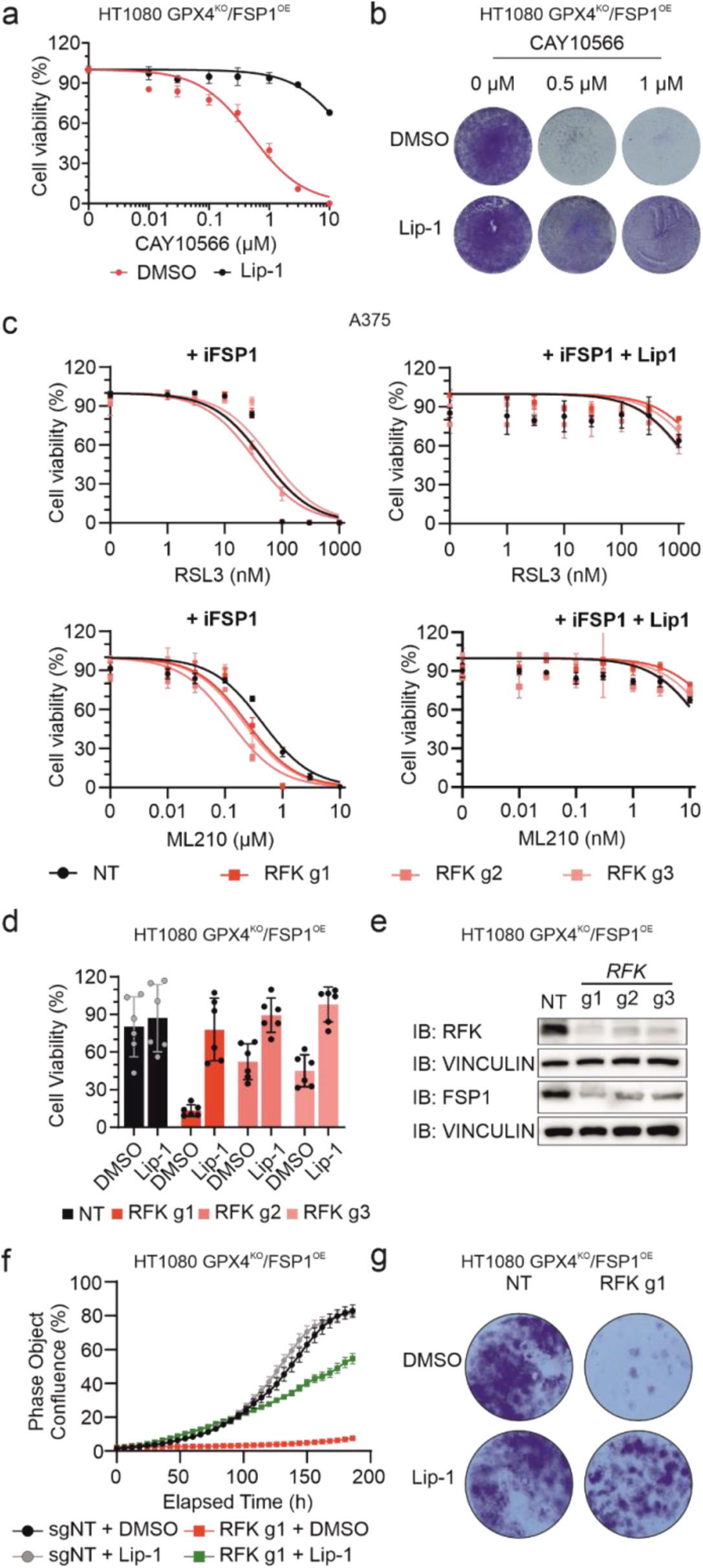
Identification of factors supporting FSP1 function. **a**, Dose***-***dependent toxicity of the SCD1 inhibitor CAY10566 in the absence or presence of the ferroptosis inhibitor Lip-1 (500 nM) in HT1080^GPX4KO/FSP1OE^ cells. Cell viability was monitored using Alamar blue after 144 h of treatment. **b**, Dose-dependent toxicity of the SCD1 inhibitor CAY10566 in the absence or presence of Lip-1 (500 nM) in HT1080 GPX4^KO^/FSP1^OE^ cells. Cell viability was monitored by crystal violet staining after 6 days of treatment. **c**, Dose-dependent toxicity of RSL3 in the presence of iFSP1 (3 µM) and/or Lip-1 (500 nM) in A375 cells transduced with either a non-targeting control (NT) or three different RFK-targeting sgRNAs. Cell viability was monitored using Alamar blue after 72 h of treatment. **d**, Cell viability of HT1080^GPX4KO/FSP1OE^ cells transduced with either a non-targeting control (NT) or three different RFK-targeting sgRNAs in the absence or presence of Lip-1 (500 nM). Cell viability was monitored using Alamar blue after 96 h. **e**, Immunoblot (IB) analysis of RFK, FSP1 and vinculin in HT1080^GPX4KO/FSP1OE^ cells transduced with either a non-targeting control (NT) or three different RFK-targeting sgRNAs. **f**, Growth curves showing the percentage of confluence over time for HT1080^GPX4KO/FSP1OE^ cells transduced with either a non-targeting control (NT) or an RFK-targeting sgRNAs in the absence or presence of Lip-1 (500 nM). Data are presented as mean ± SEM. The error was calculated per image. **g**, Cell viability of HT1080^GPX4KO/FSP1OE^ cells transduced with either a non-targeting control (NT) or an RFK-targeting sgRNAs in the absence or presence of Lip-1 (500 nM). Cell viability was monitored by crystal violet staining after 96 h.

**Extended Figure 2.**
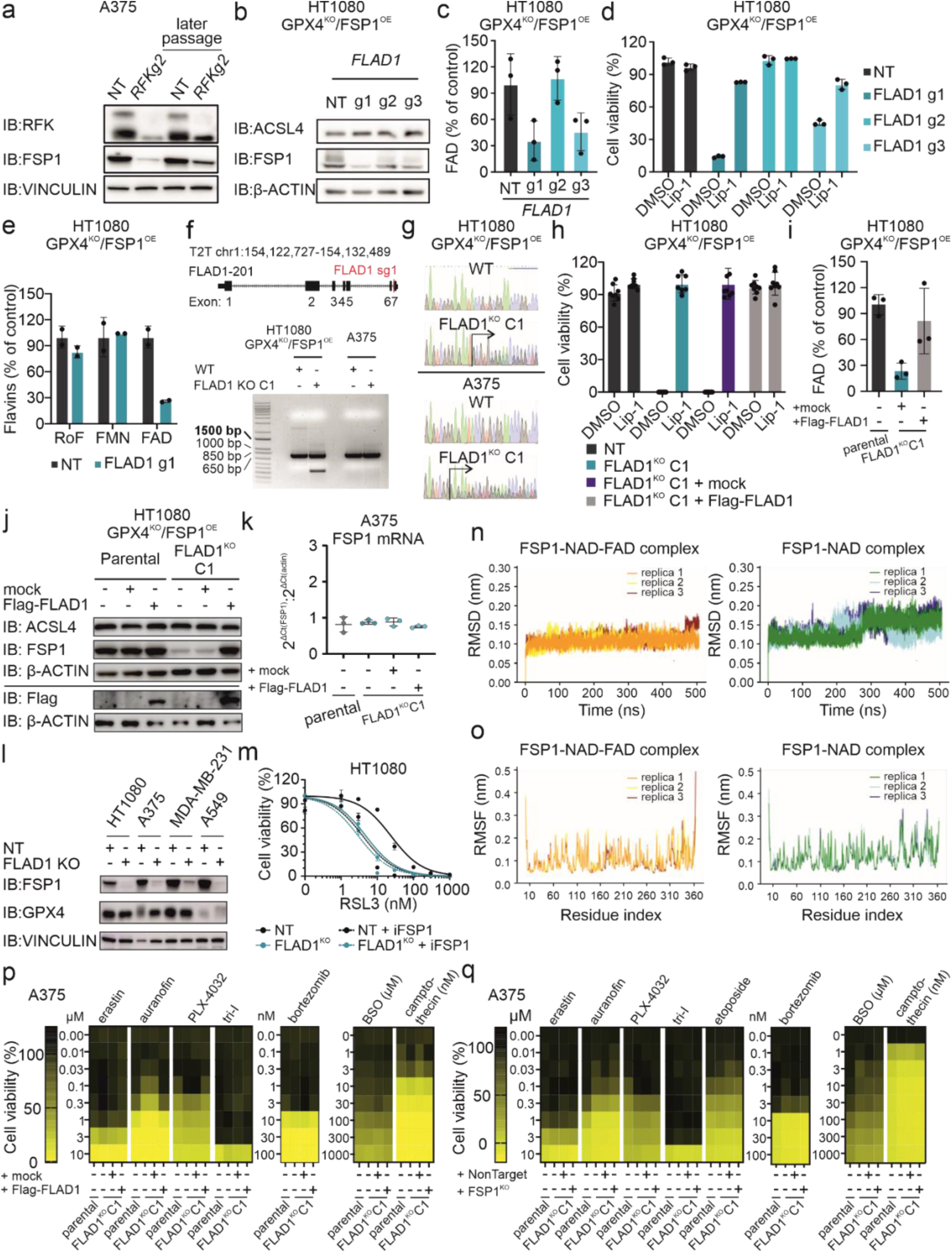
FAD deficiency disrupts FSP1 function and promotes ferroptosis susceptibility. **a**, Immunoblot (IB) analysis of RFK, FSP1 and vinculin in A375 cells transduced with either a non-targeting (NT) or RFK-targeting sgRNAs collected at an early passage or after two weeks of passaging (later passage). **b**, Immunoblot (IB) analysis of FSP1, ACSL4 and ꞵ-actin in HT1080 GPX4^KO^/FSP1^OE^ cells transduced with either a non-targeting control (NT) or three different FLAD1-targeting sgRNAs. **c**, Relative quantification of FAD in HT1080 GPX4^KO^/FSP1^OE^ cells transduced with either a non-targeting control (NT) or three different FLAD1-targeting sgRNAs. **d**, Cell viability of HT1080 GPX4^KO^/FSP1^OE^ cells transduced with either a non-targeting control (NT) or three different FLAD1-targeting sgRNAs in the absence or presence of Lip-1 (500 nM). Cell viability was monitored using Alamar blue after 96 h. **e**, Relative quantification of riboflavin (RoF), flavin mononucleotide (FMN) and flavin adenine dinucleotide (FAD) in HT1080 GPX4^KO^/FSP1^OE^ cells transduced with either a non-targeting control (NT) or an FLAD1-targeting sgRNAs (FLAD1 sgRNA1). **f**, Genotyping of wild-type (WT) and FLAD1^KO^ cells. The scheme shows the *FLAD1* gene, indicating exons and the target site of *FLAD1* sgRNA1 (red) in exon 7. The PCR products spanning the *FLAD1* sgRNA1 cut site in WT and FLAD1^KO^ (single clone, C1) from HT1080 GPX4^KO^/FSP1^OE^ and A375 cells were resolved on a 1% agarose gel. **g**, Sanger sequencing chromatograms of PCR products confirmed the presence of wild type (WT) and mutant alleles in the *FLAD1* locus in HT1080 GPX4^KO^/FSP1^OE^ and A375 cells. **h**, Cell viability of HT1080 GPX4^KO^/FSP1^OE^ non target (NT), FLAD1^KO^ single clone 1 (C1), and FLAD1^KO^ C1 cells stably overexpressing either an empty vector (mock) or Flag-FLAD1 (addback) in the absence or presence of Lip-1 (500 nM). Cell viability was monitored using Alamar blue after 24 h. **i**, Relative quantification of FAD in HT1080 GPX4^KO^/FSP1^OE^ cells “parental”, FLAD1^KO^ C1 and FLAD1^KO^ C1 cells stably overexpressing either an empty vector (mock) or Flag-FLAD1 (addback). **j**, Immunoblot (IB) analysis of ACSL4, FSP1, Flag-tag and β-actin in HT1080 GPX4^KO^/FSP1^OE^ “parental” and FLAD1^KO^ C1 cells stably overexpressing an empty vector (mock) or Flag-FLAD1 (addback). **k**, Relative mRNA levels of FSP1 measured by quantitative RT-PCR in A375 parental, FLAD1^KO^ single clone 1 (C1) and FLAD1^KO^ C1 overexpressing either an empty vector (mock) or Flag-FLAD1 (addback). **l**, Immunoblot (IB) analysis of FSP1, GPX4 and vinculin in HT1080, A375, MDA-MB-231 and A549 cells transduced with either a non-targeting (NT) or FLAD1-targeting sgRNA (FLAD1 sgRNA1). **m**, Dose-dependent toxicity of RSL3 in the absence or presence of iFSP1 (3 µM) in HT1080 cells transduced with either a non-targeting (NT) or FLAD1-targeting sgRNA (FLAD1 sgRNA1). **n**, Root mean square deviation (RMSD) of FSP1 backbone in FSP1-NAD-FAD and FSP1-NAD complexes for three 500 ns-MD simulations. **o**, Root mean square fluctuation (RMSF) of FSP1 residues in FSP1-NAD-FAD and FSP1-NAD complexes. The absence of FAD leads to more instability in the protein structure. **p**, Dose-dependent toxicity of the indicated cytotoxic compounds in A375 parental, A375 FLAD1^KO^ single clone 1 (C1) and A375 FLAD1^KO^ C1 cells stably overexpressing an empty vector (mock) or Flag-FLAD1 (addback). **q**, Dose-dependent toxicity of the indicated cytotoxic compounds in A375 parental, A375 FLAD1^KO^ single clone (C1) and A375 FLAD1^KO^ C1 transduced with either a non-targeting control (NT) or a FSP1-targeting sgRNAs (FSP1^KO^).

**Extended Figure 3.**
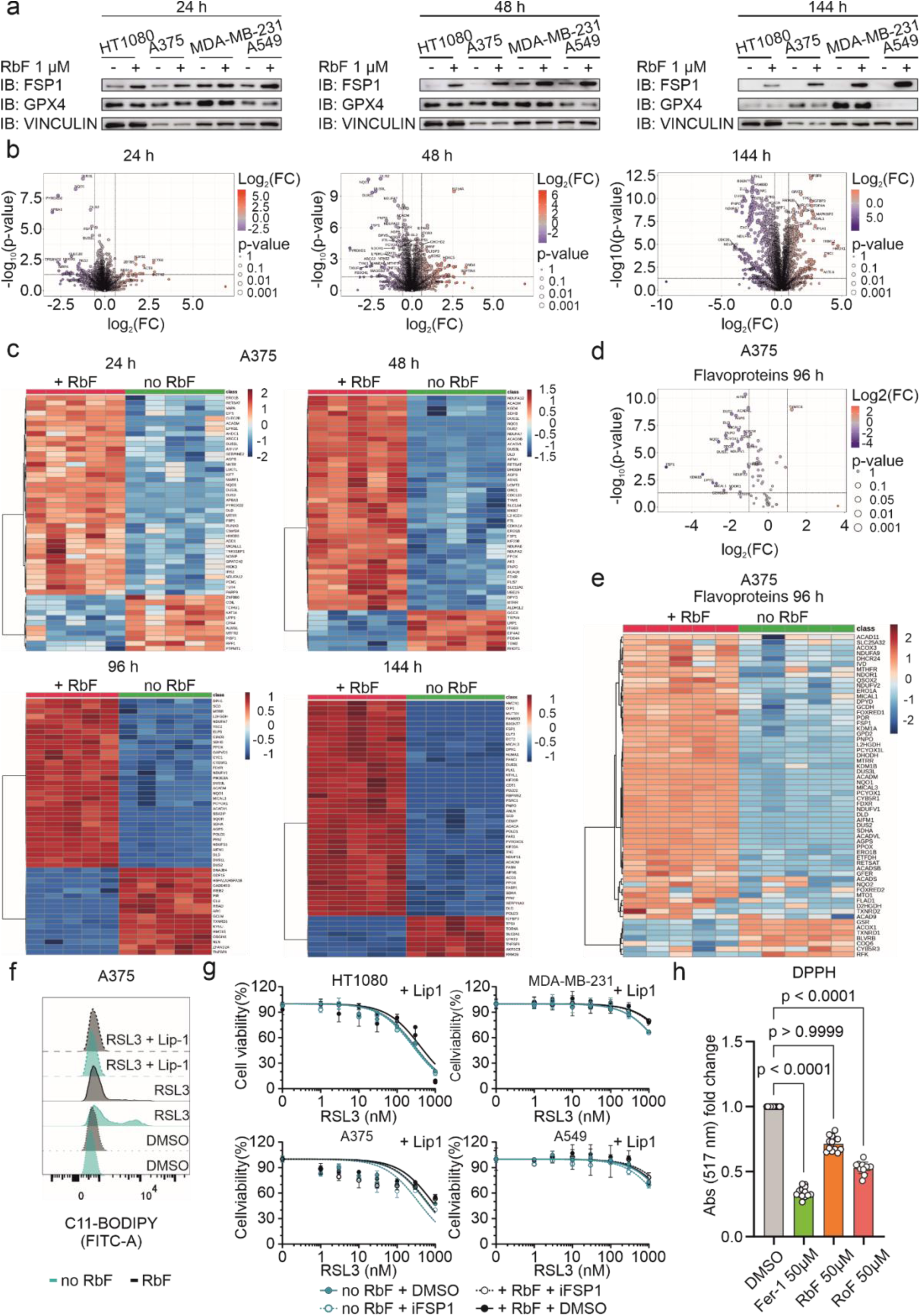
Riboflavin availability as a central determinant of ferroptosis resistance. **a**, Immunoblot (IB) analysis of FSP1, GPX4 and vinculin in HT1080, A375, MDA-MB-231 and 549 parental cell lines after 24, 48 and 144 h of growth in riboflavin-deficient medium or supplemented with 1 µM of riboflavin. **b**, Volcano plots of quantified proteins showing their change in A375 parental cells cultured in riboflavin-deficient medium for 24, 48 and 144 h. Proteins are plotted based on their fold change (FC: riboflavin deficient/normal). **c**, Heatmaps of quantified proteins showing their change in A375 parental cells cultured in riboflavin-deficient medium for 24, 48, 96 and 144 h (FC: riboflavin deficient/normal). **d**, Volcano plot of quantified flavoproteins showing their change in A375 parental cells cultured in riboflavin-deficient medium for 96 h (FC: riboflavin deficient/normal). **e**, Heatmap of quantified flavoproteins showing their change in A375 parental cells cultured in riboflavin-deficient medium for 96 h (FC: riboflavin deficient/normal). **f**, Lipid peroxidation evaluated by C11-BODIPY 581/591 staining of A375 parental cell line cultured for 72 h in riboflavin-deficient medium or supplemented with 1 µM of riboflavin and after treatment with DMSO, RSL3 (200 nM) or RSL3 (200 nM) + Lip-1 (500 nM) for 6 h. **g**, Dose-dependent toxicity of RSL3 in the presence of Lip-1 (500 nM) and iFSP1 (3 µM, when indicated) in HT1080, A375, MDA-MB-231 and A549 parental cell lines cultured in riboflavin-deficient medium or supplemented with 1 µM of riboflavin for 48 h. Cell viability was monitored using Alamar blue after 96 h of treatment. **h**, Absorbance at 517 nm corresponding to the radical initiator 2,2-diphenyl-1-picrylhydrazyl (DPPH) co-incubated with ferrostatin-1 (Fer-1), riboflavin (RbF) or roseoflavin (RoF) (n = 3). Data plotted are mean ± SD. Statistical significance was determined by one-way ANOVA followed by Dunnett’s multiple comparison test.

**Extended Figure 4.**
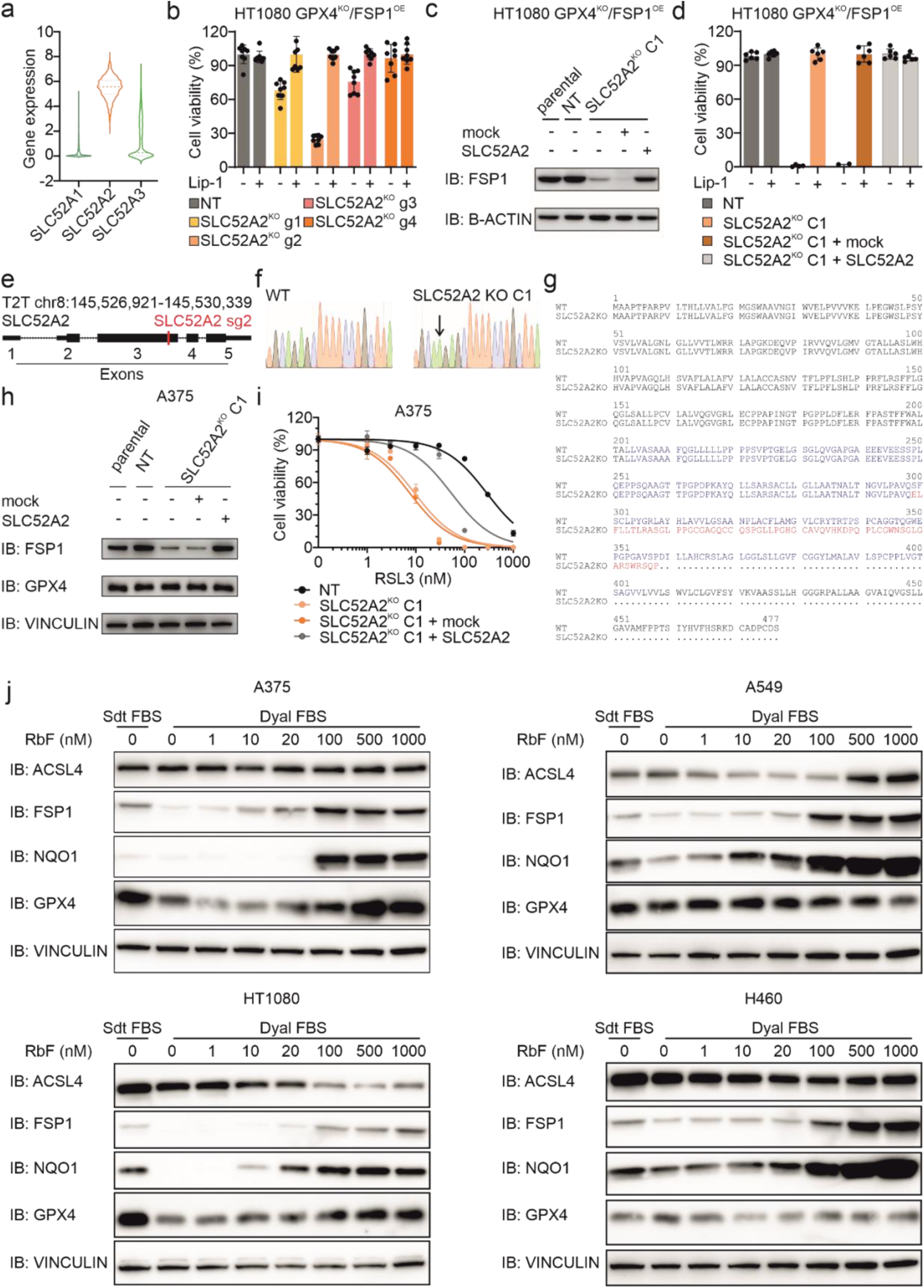
Riboflavin availability as a central determinant of ferroptosis resistance. **a**, Transcript expression levels of the riboflavin transporters SLC52A1, SLC52A2 and SLC52A3 for a panel of human cancer cell lines from Dependency Map public 23Q2 dataset (https://depmap.org/portal/, version 23Q2). **b**, Cell viability of HT1080 GPX4^KO^/FSP1^OE^ cells transduced with either a non-targeting control (NT) or four different SLC52A2-targeting sgRNAs in the absence or presence of Lip-1 (500 nM). Cell viability was monitored using Alamar blue after 96 h. **c**, Immunoblot (IB) analysis of FSP1 and ꞵ-actin in HT1080 GPX4^KO^/FSP1^OE^ “parental”, non-target control (NT), SLC52A2^KO^ single clone 1 (C1) and SLC52A2^KO^ C1 cells stably overexpressing an empty vector (mock) or SLC52A2 (addback). **d**, Cell viability of HT1080 GPX4^KO^/FSP1^OE^ cells non target control (NT), SLC52A2^KO^ single clone 1 (C1) and SLC52A2^KO^ C1 cells stably overexpressing an empty vector (mock) or SLC52A2 (addback). **e**, Genotyping of wild-type (WT) and SLC52A2^KO^ cells. The scheme shows the SLC52A2 gene, indicating exons and the target site of SLC52A2 sgRNA2 (red) in exon 3. **f**, Sanger sequencing chromatograms of PCR products confirmed the presence of WT and mutant alleles in the SLC52A2 locus in HT1080 GPX4^KO^/FSP1^OE^ and A375 cells. **g**, Schematic representation of the sequencing results obtained from the PCR product (in blue) covering the edited region (in red) in comparison with the wild-type product. **h**, Immunoblot analysis (IB) of FSP1, GPX4 and vinculin in A375 parental, non-target control (NT), SLC52A2^KO^ single clone 1 (C1) and SLC52A2^KO^ single clone 1 (C1) stably overexpressing an empty vector (mock) or SLC52A2 (addback). **i**, Dose-dependent toxicity of RSL3 in A375 non-target control (NT), SLC52A2^KO^ single clone 1 (C1) and SLC52A2^KO^ C1 stably overexpressing an empty vector (mock) or SLC52A2 (addback). **j**, Immunoblot analysis (IB) of ACSL4, FSP1, NQO1, GPX4 and vinculin in A375, A549, HT1080 and H460 parental cell lines cultured in medium supplemented with either standard FBS or dialyzed FBS and different concentrations of riboflavin (RbF).

**Extended Figure 5.**
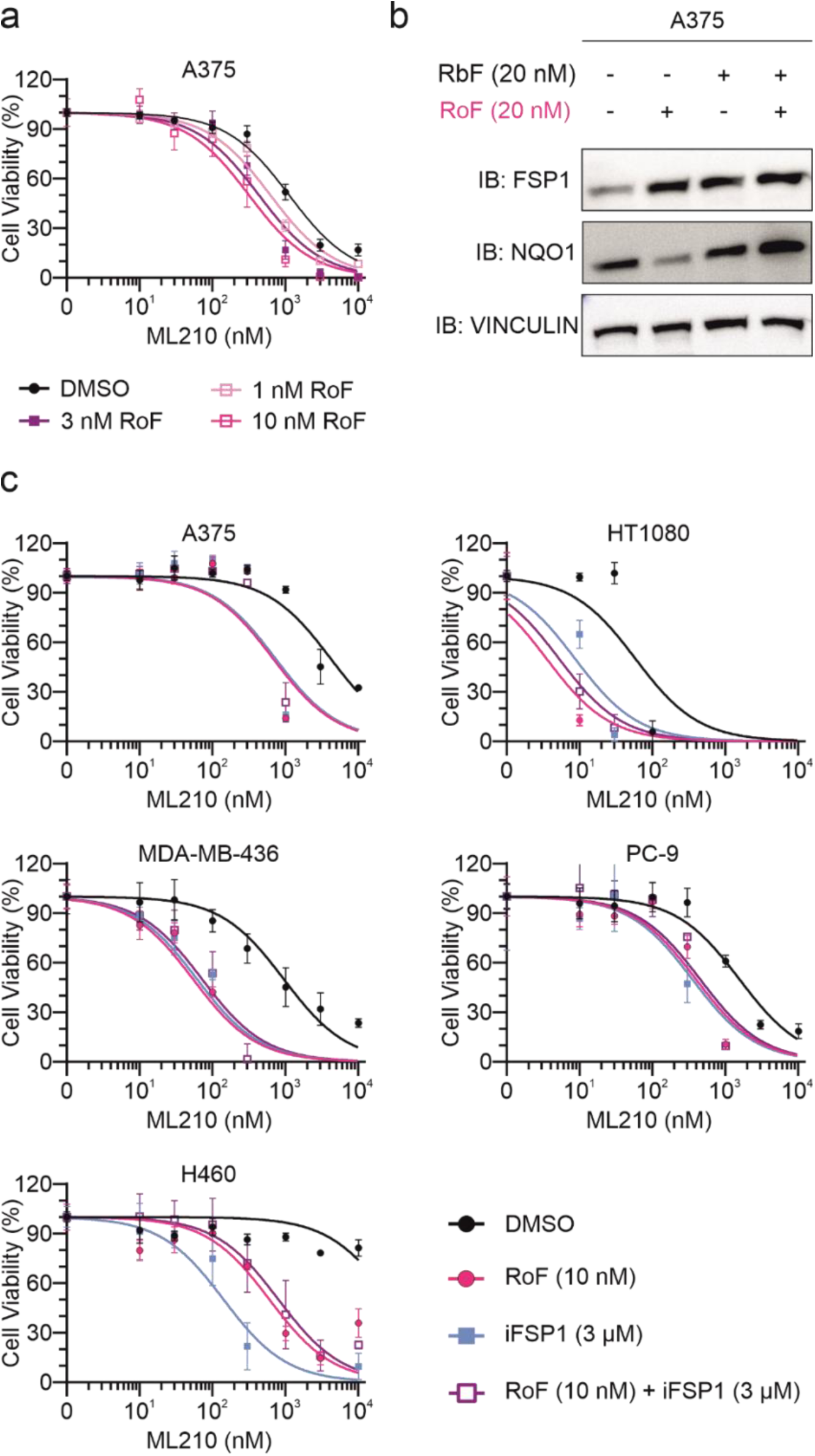
The riboflavin analogue roseoflavin disrupts FSP1 activity and promotes ferroptosis. **a**, Dose-dependent toxicity of ML210 in A375 parental cell line pre-treated with roseoflavin (RoF, 0, 1, 3 and 10 nM) for 48 h. Cell viability was monitored using Alamar blue after 48 h of treatment with ML210. Cells were cultured in low-riboflavin medium (20 nM). **b**, Immunoblot (IB) analysis of FSP1, NQO1 and vinculin in A375 parental cell line treated with roseoflavin (RoF, 20 nM) for 96 h in the absence or presence of riboflavin (RbF 20 nM). **c**, Dose-dependent toxicity of ML210 in A375, HT1080, MDA-MB-436, PC-9 and H460 parental cell lines pre-treated with roseoflavin (RoF, 10 nM) and/or iFSP1 (3 µM) for 48 h. Cell viability was monitored using Alamar blue after 48 h of treatment. Cells were cultured in low-riboflavin medium (20 nM).

## Methods

### Chemicals

Lip-1 (Sigma, cat. no. SML1414), riboflavin (Sigma, cat. no. R9504), roseoflavin (Sigma, cat. no. SML1583), RSL3 (Sigma, SML2234), ML210 (Sigma, SML0521), CAY10566 (MedChemExoress, cat. no. HY-15823), erastin (Selleckchem, cat. no. S724203), L-Buthionine-sulfoximine (BSO, Sigma cat. no. B25159), TRi-1 (MedChemExoress, cat. no. HY-125006), etoposide (MedChemExoress, cat. no. HY-13629), PLX4032 (Selleckchem, cat. no. S1267), auranofin (Sigma, cat. no. A6733), camptothecin (MedChemExoress, cat. no. HY-16560), protamine sulfate (Sigma, cat. no. P3369), bortezomib (PS-341, MedChemExpress, cat. no. HY-10227) and C11-BODIPY (581/591) (Invitrogen, cat. no. D3861) were used in this study.

### Cell lines

Human cancer cell lines HT1080, A375, MDA-MB-231, MDA-MB-436, A549 and H460 were purchased from ATCC. LOX-IMVI cells were obtained from NCI/NIH (National Cancer Institute, National Institutes of Health, USA). Cells were cultured in high-glucose DMEM GlutaMax (Gibco, cat. no. 31966-021, 4.5 g/L D-glucose) supplemented with 10% fetal bovine serum (FBS, Gibco, ref. A5256701) and 1% penicillin-streptomycin (Gibco, cat. no. 15140122). GPX4, FSP1, RFK, FLAD1, SLC52A2 knockout cells were cultured in the presence of Lip-1 (500 nM) for maintenance. All cells were cultured at 37 °C with 5% CO2 and routinely tested for Mycoplasma contamination in the laboratory.

### Lentivirus production and transduction

HEK293T cells were used to produce replication-incompetent lentiviral particles. A third-generation lentiviral packaging system consisting of transfer plasmids, envelope plasmid (pCMV-VSV-G) and packaging plasmids (pRSV_Rev and HIV-1 GAG/Pol) was used. Briefly, 700,000 cells per well were seeded on 6-well plates and cultured overnight. The next day, the cells were co-transfected with transfer, envelope and packaging plasmids (3 µg DNA/well in a proportion 1:1:1:1) using transfection reagent (X-tremeGENE HP reagent (Roche, cat. no. 06366236001)). Viral particle-containing cell culture supernatants were harvested 48 hours after transfection, filtered through a 0.45 µm membrane and used for lentiviral transduction. For transduction of the cell line of interest, 100,000 cells were seeded in a 6-well plate in medium supplemented with protamine sulfate (8 µg/mL) and Lip-1 (500 nM) and directly incubated with the lentivirus-containing supernatant for 48 hours. Transduced cells were selected by replacing the cell culture medium with fresh medium containing appropriate antibiotics, such as puromycin (1 µg/mL), blasticidin (10 µg/mL) or G418 (1 mg/mL) until non-transduced cells died.

### CRISPR-cas9 screen

HT1080 GPX4KO FSP1-FlagOE stably expressing cas9 were transduced with a lentiviral human CRISPR knockout library (VBC). This library contains 15,062 sgRNAs targeting 3007 “druggable genes” (5 sgRNAs per gene). 26 million cells were transduced to achieve 500x coverage with 1000 mg of total DNA in the presence of Lip-1 (500 nM). 48 h after infection, cells were selected with G418 (1 mg/mL) for 7 days. Once the selection was done, cells were harvested (day 0) and the remaining cells were split into two conditions (DMSO and Lip-1 500 nM) and maintained for an additional 14 days. After these 2 weeks of passaging, cells were harvested again (day 14) and genomic DNA was isolated from cell pellets (Qiagen, cat. no. 69504), followed by PCR amplification and next-generation sequencing (Illumina MiniSeq High-Output, 25M reads). The mapping of raw sequencing reads to the reference library, computing of enrichment scores and p-values were processed using the MaGeCK pipeline. MaGeCKFlute was used to visualize hits and associated pathways.

### CRISPR-cas9 mediated gene knockout

sgRNAs were selected using the VBC score (https://www.vbc-score.org/). Guides were cloned using annealed oligonucleotides (Eurofins genomics) with specific overhangs complementary to BsmBI-digested lentiCRISPRv2-puro or lentiCRISPRv2-blast (Addgene, cat. no. 98290 and 98293, respectively) and selected for 4-7 days. Knockout efficiency was validated by immunoblotting (when antibodies were available) and sequencing of genomic DNA. In the case of the FLAD1 gene, the knockout was also confirmed by measuring FAD levels. Cells were used as pools unless stated otherwise. The sequences of the guides used in the study are provided below.

hGPX4_sgRNA1 – GAAGCGCTACGGACCCATGG

hFSP1_sgRNA1 – GGTGCAGAGAATCACCAGGT

hRFK_sgRNA1 – GAGATGTCCATAAGATGG

hRFK_sgRNA2 – ACTTGACCCCGGCAGAAGTA

hRFK_sgRNA3 – CTATGGGGAAATCCTCAATG

hFLAD1_sgRNA1 - AGTCGGGAGAATACCGTG

hFLAD1_sgRNA2 – GATCTGTGCCATAATGC

hFLAD1_sgRNA3 – GAGTAGGGGTCAGTCCGG

hSLC52A2_sgRNA1 – GCCAGGAAGCAGGCCAG

hSLC52A2_sgRNA2 - TAAGCAGGAAAAGCTCTGCA

hSLC52A2_sgRNA3 – CTGGCTGCCACCTTCACGT

hSLC52A2_sgRNA4 - GTGGGTCAGCACCGGAC

### Stable protein overexpression

HT1080 and A375 cells overexpressing human FSP1, FLAD1 and SLC52A2 proteins were generated by transduction using lentivirus particles as described above. For cloning FSP1, FLAD1 and SLC52A2 expression vectors, the sequences (CCDS7297.1, CCDS1078.1 and CCDS6423.1, respectively) were codon-optimized (IDT Codon Optimization Tool) and synthesized by IDT as gBlocksTM and cloned using ClonExpress II One Step Cloning Kit (Vazyme, cat. no. C112) into p442-IRES-blast or p442-IRES-neo vectors. Afterward, the plasmids were delivered into the cell of interest using lentivirus particles as described above.

### Cell viability assays

Cells were seeded at 1000 cells/well on 96-well plates and after 4-6 hours treated with the following compounds: RSL3, ML210, iFSP1, Lip-1, erastin, BSO, TRi-1, etoposide, PLX-4032, auranofin, camptothecin, bortezomib and roseoflavin. Cell viability was assessed 72-96 hours after treatment using the Alamar Blue method as an indicator of viable cells. Alamar Blue solution was made by dissolving 1 g of resazurin sodium salt in 100 mL of PBS and filtered through a 0.22 µm membrane. The working solution was made fresh by adding 80 µL of the stock solution to every 10 mL of DMEM medium. After 3 hours of incubation, viability was estimated by measuring fluorescence using a 540/35 excitation filter and a 590/20 emission on a Spark® microplate reader (Tecan, Zürich, Switzerland).

### Live-Cell Imaging and Proliferation analysis

Cell proliferation was monitored using the IncuCyte® Live-Cell Analysis System S3/SX1 (Sartorius). The plate was placed in the IncuCyte incubator, and phase-contrast images were acquired every 6 hours for a total duration of 5-7 days. Image acquisition was performed at 10x magnification, and confluence was determined using the IncuCyte integrated analysis software.

### Cell lysis and immunoblotting

Cells were lysed in RIPA buffer containing protease inhibitor cocktail (Roche, cat. no. 11697498001) and centrifuged at 20,000g for 30 minutes at 4 °C. Protein concentration was normalized by the bicinchoninic acid (BCA) assay (Thermo Fisher, cat. no. 23235), and samples were heated at 70 or 90 °C for 10 minutes in the presence of SDS-containing loading buffer. Proteins (20 µg per lane) were resolved on 12% SDS-PAGE gels and subsequently electroblotted onto nitrocellulose or PVDF membranes. The membranes were blocked in non-fat milk 5% in TBT-T (20 mM Tris-HCl, 150 mM NaCl and 0.1% Tween-20) for 1 hour at RT. Subsequently were incubated overnight at 4 °C with primary antibodies diluted with 5% milk or bovine serum albumin (BSA) in TBS-T against FSP1 (1:10, antibody raised against recombinant human FSP1 protein, clone 6D8-11, developed in Helmholtz Zentrum München), GPX4 (1:1,000, Abcam, cat. no. ab125066 or 1:500, Proteintech, cat. no. 67763-1-Ig), ACSL4 (1:500, Santa Cruz, cat. no. sc-271800), FLAD1 (1:250, Santa Cruz, cat. no. sc-376819), RFK (1:250, Santa Cruz, cat. no. sc-398830 or 1:500, Abbexa cat. no. abx124688), NQO1 (1:5,000, Santa Cruz, cat. no. sc-32793), Flag-tag (1:1,000, Sigma-Aldrich, cat. no. F3165), β-actin (1:5,000, Sigma-Aldrich, cat. no. A5441) and vinculin (1:1,000, Santa Cruz, sc-73614). The next day, membranes were washed with TBS-T and incubated with horseradish peroxidase-labeled secondary antibodies (1:3,000, Cell Signaling, cat. no. 7074, 7076 and 7077) diluted in non-fat milk 5% in TBS-T for 2 hours at RT. Finally, membranes were washed, and antibody-antigen complexes were detected by chemiluminescence using enhanced chemiluminescence substrate (BioRad, cat. no. 107-5061) on Amersham ImageQuant 800 (Cytiva).

### Genotyping

FLAD1 and SLC52A2 single-clone knockout cells were confirmed through genotyping.

Genomic DNA from WT and knockout cell pellets was extracted by proteinase K digestion. Subsequently, the sgRNA-targeting regions were amplified by PCR using primers flanking the sgRNA cut-site and analyzed via Sanger sequencing to identify WT and knockout alleles sequences. Indels were characterized using CRISP-ID web-based application^1^. To further confirm the precise editing events at the FLAD1 locus in the knockout cells, the PCR product was cloned into a pJET1.2 cloning vector using CloneJET PCR Cloning Kit (Thermo Fisher, cat. no. K1231), followed by PCR amplification of 5 colonies and Sanger sequencing analysis. The sequences of primers used are provided below.

FLAD1_Fw – CTCCACCCTTGCACTAGAGG

FLAD1_Rv – CCCCTTAGAGTGAGCACAGC

SLC52A2_Fw – GACTTCCTTGAGCGTTTTCCC

SLC52A2_Rv - GGGGAACATAGCAACAGCG

Preparation of riboflavin-deficient medium

The DMEM medium without riboflavin was purchased from PAN biotech (cat. no. P04-03584) and supplemented with GlutaMax (Gibco, cat. no. 35050-061), 10% dialyzed fetal bovine serum (FBS, Gibco, ref. A5256701), 1% penicillin-streptomycin (Gibco, cat. no. 15140122), 15 mg/L phenol red (Sigma, cat. no. P0290) and riboflavin (1 µM, unless noted otherwise) as needed for comparison. Riboflavin stock solution (20 mg/L) was freshly prepared by dissolving riboflavin (Sigma, cat. no. R9504) in PBS and sterilizing through a 0.22 µm membrane filter.

### Dialysis of Fetal Bovine Serum

Fetal bovine serum (FBS) was dialyzed to remove small molecules, including riboflavin, using tubing cellulose membranes (Sigma, cat. no. D9527) with a 14 kDa molecular weight cutoff. Dialysis was performed against PBS at a 1:10 (v/v) ratio, with the buffer replaced every 24 h over five consecutive cycles under constant magnetic stirring at 4 °C. The dialyzed serum was subsequently filtered through a 0.22 µm membrane for sterility, aseptically transferred into sterile containers, aliquoted, and stored at −20 °C until further use.

### Cell viability assays in riboflavin-deficient medium

HT1080, A375, MDA-MB-231 and A549 parental cell lines were pre-incubated in riboflavin-deficient medium for 48 hours, seeded at 1,000 cells/well in 96-well plates and after 4-6 hours treated with RSL3 (0-1000 nM) in the absence or presence of iFSP1 (3 µM) and/or Lip-1 (500 nM). Cell viability was assessed after 96 hours of treatment using the Alamar Blue method.

To evaluate the effect of roseoflavin, HT1080, A375, MDA-MB-231, A549, MDA-MB-436, H460, LOX-IMVI and PC-9 parental cell lines were cultured in low-riboflavin medium (20 nM). 3,000 cells/well were seeded in 96-well plates and after 4 – 6 hours pre-treated with roseoflavin (0, 1, 3 and 10 nM) for 48 hours. Following pre-treatment with roseoflavin, cells were treated with ML210 (0-10 µM) or iFSP1 (3 µM) for an additional 48 hours, after which cell viability was assessed using the Alamar Blue method.

### Riboflavin, FMN and FAD measurements

Water soluble metabolite measurements were made by liquid chromatography-mass spectrometry (LC/MS) analysis. For sample preparation, 1 million cells were harvested, washed with PBS and rapidly frozen with liquid nitrogen.

Water-soluble metabolites were extracted with 500 µL of ice-cold MeOH/H2O (80/20, v/v) containing 0.01 μM lamivudine and 1 µM each of D2-glucose, D4-succinate, D5-glycine and 15N-glutamate (Sigma-Aldrich). After centrifugation of the resulting supernatants were evaporated in a rotary evaporator (Savant, Thermo Fisher Scientific, Waltham, USA). Dry sample extracts were redissolved in 150 μL 5 mM NH4OAc in CH3CN/H2O (50/50, v/v). 20 µL supernatant was transferred to LC-vials. Metabolites were analyzed by LC-MS using the following settings: 3 μL of each sample was applied to a XBridge Premier BEH Amide (2.5 μm particles, 100 × 2.1 mm) UPLC-column (Waters, Dublin, Ireland). Metabolites were separated with Solvent A, consisting of 5 mM NH4OAc in CH3CN/H2O (40/60, v/v) and solvent B consisting of 5 mM NH4OAc in CH3CN/H2O (95/5, v/v) at a flow rate of 200 µL/min at 45 °C by LC using a DIONEX Ultimate 3000 UHPLC system (Thermo Fisher Scientific, Bremen, Germany). A linear gradient starting after 2 min with 100 % solvent B decreasing to 10% solvent B within 23 min, followed by 16 min 10% solvent B and a linear increase to 100% solvent B in 2 min was applied. Recalibration of the column was achieved by a 7-minute pre-run with 100% solvent B before each injection.

All MS analyses were performed on a high-resolution Q Exactive mass spectrometer equipped with a HESI probe (Thermo Fisher Scientific, Bremen, Germany) in alternating positive- and negative-full MS mode with a scan range of 69.0-1000 m/z at 70K resolution and the following ESI source parameters: sheath gas: 30, auxiliary gas: 1, sweep gas: 0, aux gas heater temperature: 120 °C, spray voltage: 3 kV, capillary temperature: 320 °C, S-lens RF level: 50. XIC generation and signal quantitation was performed using TraceFinder™ V 5.1 (Thermo Fisher Scientific, Bremen, Germany) integrating peaks which corresponded to the calculated monoisotopic metabolite masses (MIM +/-H+ ± 3 mMU). All analyses were performed in three independent biological replicates.

### RNA extraction, cDNA synthesis and RT-qPCR

Isolation of total RNA from cell pellets was performed using TRIzol reagent (Invitrogen, cat. no. 15596026) according to the manufacturer instructions. cDNA synthesis was done using HiScript III 1st Strand cDNA Synthesis Kit (Vazyme, cat. no. R312) and hexamer primers in accordance to the manufacturer’s protocol. RT-qPCR was performed and analysed with a Mastercycler ep realplex (Eppendorf) or CFX Connect (Biorad) using SYBR Green reagent. Gene expression was normalized to ACTB as a housekeeping gene using the ΔΔct method.

The qPCR primer pairs are indicated below.

hFSP1_1_Fw – GACTCCTTCCACCACAATGTGG

hFSP1_1_Rv – CAGCACCATCTGGTTCTTCAGG

hFSP1_2_Fw – CCGCTATCCAGGCCTATGAG

hFSP1_2_Rv - AATCTCTGCTGCCATCTCCA

Lipid peroxidation assays using C11-BODIPY (581/591)

50,000 cells per well were seeded on 6-well plates one day prior to the experiment. The next day, cells were treated with the following conditions (i) DMSO, (ii) RSL3 200 nM and (iii) RSL3 200 nM + Lip-1 500 nM for 6 hours. After treatment, cells were washed with PBS and incubated with C11-BODIPY (1 µM) for 30 minutes at 37 °C before they were harvested by trypsinization. Subsequently, cells were resuspended in 400 µL of PBS supplemented with 2% FBS followed by analysis using a flow cytometer (FACS Canto II, BD Biosciences). Data was collected from the FITC detector (for the oxidized form of BODIPY) with a 502LP and 530/30 BP filter and from the PE detector (for the reduced form of BODIPY) with 556 LP and 585/42 BP filter. At least 10,000 events were analyzed per sample. Data was analyzed using FlowJo Software. The ratio FITC/PE (oxidized/reduced ratio) was calculated as follows: (median FITC-A fluorescence – median FITC-A fluorescence of unstained samples)/(median PE-A fluorescence – median PE-A fluorescence of unstained samples).

For riboflavin deprivation experiments, cells were pre-incubated in riboflavin-deficient medium or medium supplemented with riboflavin (1 µM) for 72 hours before undergoing the treatments described above.

### Epilipidomics analysis

500,000 A375 parental cells were seeded on 15-cm dishes in a standard DMEM medium. After 4 hours, cells were divided into two conditions: (i) riboflavin-deficient medium and (ii) supplemented with riboflavin (1 µM) medium. Cells were maintained under these conditions for 72 hours. Following the 72-hour incubation, cells were treated with one of the following conditions for 6 hours: (i) DMSO, (ii) RSL3 200 nM, and (iii) RSL3 200 nM + Lip-1 500 nM. After treatment, cells were harvested and washed with PBS. The cell pellets were snap-frozen in liquid nitrogen and stored at −80 °C until further analysis.

Lipids were extracted according to Folch protocol^2^. All solvents contained 1 μg/mL butylated hydroxytoluene and were cooled on ice before use. Briefly, cell pellets (3-5 x106 cells) collected in PBS were washed, centrifuged, and resuspended in 50 μL of H2O in 2 mL Eppendorf tubes. 5 µL of SPLASH® LIPIDOMIX® (Avanti Polar Lipids Inc., Alabaster, AL, USA) were added, tubes were vortexed for 10 s and left on ice for 15 min. 365 μL ice-cold MeOH was added, the samples were vortexed for 10 s, then 740 μL of ice-cold CHCl3 were added, vortexed for 10 s, and the samples were incubated for 1 h in a rotary shaker (40 rpm) at 4 °C. Phase separation was induced by adding 225 μL of H2O, the samples were vortexed for 10 s and centrifuged at 2000g for 10 min. 770 μL of the lower phase was transferred to 1.5 mL Eppendorf tubes and dried in a vacuum concentrator at 40 mbar and 20 °C. Meanwhile, lipids from the remaining upper phase were re-extracted by adding 400 μL of CHCl3:MeOH 2:1 (v/v), vortexed for 10 s, 100 μL H2O was added to promote phase separation, vortexed for 10 s, centrifuged at 2000g for 10 min, and 380 μL of the lower phase was transferred to the 1.5 mL Eppendorf tube containing the first extract portion and the sample was continued to dry in the vacuum concentrator.

The dried lipid extracts were reconstituted in 125 μL of i-PrOH, centrifuged at 10000g for 5 min, and 120 μL were transferred into vials with glass inserts for LC-MS analysis. The upper phase remained after extraction was dried in the vacuum concentrator at 20 mbar and 20 °C and used for total protein quantification. Group-specific samples (gQC) were prepared by pooling 40 μL of the corresponding individual samples, total quality control (tQC) samples were prepared by mixing 40 μL of gQC samples.

Reversed phase liquid chromatography (RPLC) was carried out on a Vanquish Horizon (Thermo Fisher Scientific, Bremen, Germany) equipped with an Accucore C30 column (150 x 2.1 mm; 2.6 µm, 150 Å, Thermo Fisher Scientific, Bremen, Germany). Lipids were separated by gradient elution with solvent A (MeOH/H2O, 1:1, v/v) and B (i-PrOH/MeCN/H2O, 85:15:5, v/v/v) both containing 5 mM HCOONH4 and 0.1% (v/v) HCOOH. Separation was performed at 50 °C with a flow rate of 0.3 mL/min using the following gradient: 0-20 min – 10 to 86 % B (curve 4), 20-22 min – 86 to 95 % B (curve 5), 22-26 min – 95 % isocratic, 26-26.1 min – 95 to 10 % B (curve 5) followed by 5 min re-equilibration at 10% B.

Mass spectrometry was performed on Thermo Scientific Orbitrap Exploris 240 (Thermo Fisher Scientific) equipped with a heated electrospray ionization (HESI) source with an EASY-IC unit for lock mass correction and operated with the following global HESI parameters: sheath gas 40 arbitrary units, auxiliary gas 10 arbitrary units, sweep gas 1 arbitrary units, spray voltage 3.5 kV (positive mode) or 2.5 kV (negative mode), ion transfer tube temperature 300 °C, vaporizer temperature 370 °C, S-lens RF level 35%, EASY-IC lock mass correction was set to RunStart. For identification of oxidized lipids, 6 μL of gQC samples were applied onto the LC column and MS data was recorded in ionization polarity switch mode (0-22.5 min – negative, 22.5-39.9 min – positive) with the instrument operating in semi-targeted data dependent acquisition (stDDA) mode with 6 MS/MS scans per cycle. Full scans (MS1) had the following settings: Orbitrap resolution 60000 at m/z 200, scan range m/z 500-980 (negative mode) and 480-1000 (positive mode), absolute AGC value set to standard, maximum injection time set to auto, 1 microscan. StDDA MS2 scans had the following settings: 1.5 m/z precursor selection isolation window, Orbitrap resolution 30000 at m/z 200, stepped higher-energy collisional dissociation (HCD) at normalized collision energies (nCE) of 22-32-43% (negative mode) or 32-43-54% (positive mode), absolute AGC value 1.105, maximum injection time 200 ms, 2 microscans. The following filters were applied prior to MS2 scans: dynamic exclusion after 5 times if occuring within 6 s, exclusion duration 5 s, mass tolerance ±5 ppm, isotope exclusion; allowed charge state 1; targeted mass inclusion using the mass list of in silico predicted oxidized lipids (667 individual m/z values split into 3 LC-MS runs), inclusion mass tolerance ±5 ppm. Data was acquired in profile mode.

Oxidized lipids were identified using LPPtiger2 software (Fedorova Lab)^3,4^. For relative quantification of oxidized lipids identified in stDDA runs, retention time-scheduled parallel reaction monitoring (PRM) in ionization polarity switch mode (0-22 min – negative, 22-40 min – positive) at the resolution of 15000 at m/z 200, absolute AGC value of 1.105 and a maximum injection time of 100 ms. The isolation window for precursor selection was 1.5 m/z. HCD nCE for every target was chosen based on optimization PRM runs of gQC samples (fixed nCE from 15 to 40%). Data was acquired in profile mode. PRM data were processed in Skyline v. 24.1.0.199 (MacCoss Lab)5 considering fragment anions of oxidized fatty acyl chains as quantifier. The obtained peak areas were normalized by appropriate lipid species from SPLASH® LIPIDOMIX® Mass Spec Standard (Avanti), e.g. by LPC(18:1(d7)), LPE(18:1(d7)), PC(15:0/18:1(d7)), or PE(15:0/18:1(d7)), and protein concentration measured for the corresponding sample. Normalized peak areas were further log-transformed and autoscaled in MetaboAnalyst online platform (https://www.metaboanalyst.ca, Xia Lab)^6^.

### Proteomic analysis

A375 FLAD1KO clone 1 (C1) and FLAD1KO C1 Flag-FLAD1OE (addback) cells were seeded at a density of 1 million cells on 10-cm dishes. After 24 hours, cells were harvested, washed once with PBS, snap-frozen in liquid nitrogen and stored at −80 °C until further processing.

In a separate experiment, A375 parental cells were cultured in DMEM medium either deficient in riboflavin or supplemented with 1 µM of riboflavin. Cells were seeded at varying densities 500,000, 250,000, 60,000 and 15,000 cells per 10-cm dish to collect at different time points: 24, 48, 96 and 144 hours, respectively. At each time point, 1 million cells were harvested, washed with PBS, snap-frozen in liquid nitrogen and stored at −80 °C until further analysis.

For proteomics analysis, cell pellets were lysed in RIPA buffer supplemented with protease and phosphatase inhibitor cocktails (Roche, cat. no. 11697498001 and 4906845001, respectively) and 1% (v/v) benzonase (Merck, cat. no. E1014). Lysates were sonicated with 3 cycles of 10 seconds on/30 seconds off at 40% amplitude (Bioruptor Plus). Removal of residual cell debris was made by centrifugation at 17,000g for 2 hours at 4 °C. The supernatants containing soluble proteins were collected, and protein concentrations were determined using the BCA assay (Thermo Fisher, cat. no. 23235). Protein concentrations were normalized to 1 µg/µL using water. Normalized protein samples were aliquoted and stored at −80 °C until further analysis.

Tryptic digestion of proteins (Input: 10 µg) was performed using an AssayMAP Bravo liquid handling system (Agilent technologies) running the autoSP3 protocol according to Müller et al.7. The resulting peptides were vacuum dried and stored at −20 °C until LC-MS/MS analysis.

The LC-MS/MS analysis was carried out on an Ultimate 3000 UPLC system directly connected to an Orbitrap Exploris 480 mass spectrometer (both Thermo Fisher Scientific) for a total of 120 min injecting an equivalent of 1 µg of peptide. Peptides were online desalted on a trapping cartridge (Acclaim PepMap300 C18, 5 µm, 300 Å wide pore; Thermo Fisher Scientific) for 3 min using 30 µl/min flow of 0.05% TFA in water. The analytical multistep gradient (300 nl/min) was performed using a nanoEase MZ Peptide analytical column (300 Å, 1.7 µm, 75 µm x 200 mm, Waters) using solvent A (0.1% formic acid in water) and solvent B (0.1% formic acid in acetonitrile). For 102 min the concentration of B was linearly ramped from 4% to 30%, followed by a quick ramp to 78%, after two minutes the concentration of B was lowered to 2% and a 10 min equilibration step appended. Eluting peptides were analyzed in the mass spectrometer using data independent acquisition (DIA) mode. A full scan at 120k resolution (380-1400 m/z, 300% AGC target, 45 ms maxIT) was followed by 47 MS2 windows covering the mass range from 400-1000 m/z with variable width and overlapping by 1 Da (30k resolution, AGC target 1000%, maxIT 54 ms, 28% HCD collision energy). Each sample was followed by a wash run (40 min) to minimize carry-over between samples. Instrument performance throughout the course of the measurement was monitored by regular (approx. one per 48 hours) injections of a standard sample and an in-house shiny application.

All sample handling (sample preparation and LC-MS/MS analysis) have been performed in a block randomization order8.

Analysis of DIA RAW files was performed with Spectronaut (Biognosys, version 19.1.240724.62635; Bruderer R. et al. (2015)^9^ in directDIA+ (deep) library-free mode. Default settings were applied with the following adaptions. Within DIA Analysis under Identification the Precursor PEP Cutoff was set to 0.01, the Protein Qvalue Cutoff (Run) set to 0.01 and the Protein PEP Cutoff set to 0.01. In Quantification the Proteotypicity Filter was set to Only Protein Group Specific, the Protein LFQ Method was set to MaxLFQ and the Quantification window was set to Not Synchronized (SN 17). The data was searched against the human proteome from Uniprot (human reference database with one protein sequence per gene, containing 20,597 unique entries from ninth of February 2024) and the contaminants FASTA from MaxQuant (246 unique entries from twenty-second of December 2022). Conditions were included in the setup.

Statistical Analysis of Proteomics Data

Statistical analysis was performed using MetaboAnalyst software10, with Spectronaut output files as input. After Spectronaut analysis, the data set comprised the identified proteins, along with LFQ intensities. The data were filtered to remove features matched with contaminant or reverse sequences. The missing values were replaced with LoDs (1/5 of minimum positive values of corresponding variables) and data were log10 transformed and used for further statistical analysis. The fold change threshold was taken to be 1.5, and RAW P value cutoff was 0.05 to get the DEPs. Volcanoplot and heatmap representing the expression of top DEPs was also obtained from statistical analysis in MetaboAnalyst. Metascape tool^11^ was used to obtain statistically enriched terms (GO/Kyoto Encyclopedia of Genes and Genomes pathway [KEGG] terms, canonical pathways) for all DEPs. All analyses were performed in five independent biological replicates.

### DPPH Assay

To assess whether riboflavin and roseoflavin act as radical-trapping antioxidants, the compounds were tested in a cell-free antioxidation assay. 10 mM compound was diluted in 1 mL 2,2-diphenyl-1-picrylhydrazyl (DPPH, 0.05 mM in methanol, Sigma-Aldrich) to a final concentration of 50 µM. Samples were rotated at room temperature for 10 minutes before they were transferred into a clear 96-well plate in quadruplicates. Absorbance was measured at 517 nm with an EnVision 2104 Multilabel plate reader (PerkinElmer). Experiment was performed in three biological replicates, each consisting of four technical replicates. DMSO was used as normalization control; Ferrostatin-1 was used as a positive control potent antioxidant.

### Molecular dynamics simulations

The crystal structure of the human FSP1 complex (PDB ID: 8WIK)^12^ was used as the initial structure to prepare two systems: the FSP1-NAD complex and the FSP1-NAD-FAD complex. The structure of FAD was obtained by substituting OH from C6 with H in the 6FA ligand present in PDB ID 8WIK. The protonation states of the residues at pH 7.4 were determined using the protein preparation wizard tool from Maestro (Schrödinger v.2024.2)^13,14^. Ligand parameters for NAD+ were obtained from the Manchester Amber parameter database^15^. The partial charges of FAD were determined using AmberTools23^16^ by the restrained electrostatic potential charges (RESP) method. A quantum mechanical calculation was performed to obtain the electrostatic potentials using Gaussian 09^17^, Hartree Fock, the 6-31G* basis set and full optimisation. The bond, angle, torsion, and van der Waals parameters were generated using the general AMBER force field (GAFF)^18^ for co-factors, and the AMBER-ILDN force field^19^ for the protein.

Three independent simulations of each system, the FSP1-NAD complex and the FSP1-NAD-FAD complex, were carried out using GROMACS 2024.2 (Abraham et al. GROMACS 2024.2)^20^. Each system was solvated with TIP3P^21^ water molecules with a margin of at least 10 Å, and Na+ and Cl− ions were added to ensure system neutrality at an ion concentration of 150 mM. The steepest descent method was used to perform energy minimization for 5000 steps on each system using positional restraints of 1000 kJ/mol/nm2 on the heavy atoms of the protein and cofactors. The systems were then equilibrated with an NVT ensemble to achieve constant temperature at 300 K using the velocity rescaling thermostat^22^. Next, the systems were equilibrated to a constant pressure of 1 bar using the cell rescaling barostat^23^. Following equilibration, positional restraints were gradually lifted from the heavy atoms of the protein and the cofactors (500,100,10 kJ/mol/nm2). Production runs of 500 ns were performed with a timestep of 2 fs. The temperature coupling was achieved with the velocity rescaling at time constant of 0.1 ps and pressure coupling was achieved with cell rescaling at compressibility of 4.5.10−5 bar−1 and a time constant of 5 ps. LINCS^24^ algorithm was used to constrain covalent H-bonds and SETTLE25 algorithm was used to constrain the solvent bond lengths. The particle mesh Ewald (PME)^26,27^ method was used to calculate the electrostatic forces with a real-space cutoff of 1.2 nm, PME order of four, and a Fourier grid spacing of 1.2 Å. A cut-off of 1.2 nm was used for the calculation of the Van der Waals interactions. Data analysis (root mean square deviation, root mean square fluctuation and DSSP analysis) was performed using GROMACS 2024.2 (Abraham et al. GROMACS 2024.2)^20^.

### Human cancer cell line datasets

Human cancer cell line datasets were obtained from the DepMap portal (https://depmap.org/portal/, version 23Q2).

### Statistical analysis

All experiments (except those described otherwise in the legend) were performed independently at least twice. Graphs were generated using GraphPad Prism v10 (GraphPad Software) if not stated otherwise.

## Reference

1 Xu, L., Davis, T. A. & Porter, N. A. Rate constants for peroxidation of polyunsaturated fatty acids and sterols in solution and in liposomes. J Am Chem Soc 131, 13037–13044 (2009). 10.1021/ja9029076

2 Stockwell, B. R. Ferroptosis turns 10: Emerging mechanisms, physiological functions, and therapeutic applications. Cell 185, 2401–2421 (2022). 10.1016/j.cell.2022.06.003

3 Yang, W. S. et al. Regulation of ferroptotic cancer cell death by GPX4. Cell 156, 317–331 (2014). 10.1016/j.cell.2013.12.010

4 Friedmann Angeli, J. P., et al. Inactivation of the ferroptosis regulator Gpx4 triggers acute renal failure in mice. Nat Cell Biol 16, 1180–1191 (2014). 10.1038/ncb3064

5 Angeli, J. P. F., Shah, R., Pratt, D. A. & Conrad, M. Ferroptosis Inhibition: Mechanisms and Opportunities. Trends Pharmacol Sci 38, 489–498 (2017). 10.1016/j.tips.2017.02.005

6 Doll, S. et al. FSP1 is a glutathione-independent ferroptosis suppressor. Nature 575, 693–698 (2019). 10.1038/s41586-019-1707-0

7 Bersuker, K. et al. The CoQ oxidoreductase FSP1 acts parallel to GPX4 to inhibit ferroptosis. Nature 575, 688–692 (2019). 10.1038/s41586-019-1705-2

8 Mishima, E. et al. A non-canonical vitamin K cycle is a potent ferroptosis suppressor. Nature 608, 778–783 (2022). 10.1038/s41586-022-05022-3

9 Peck, B. et al. Inhibition of fatty acid desaturation is detrimental to cancer cell survival in metabolically compromised environments. Cancer Metab 4, 6 (2016). 10.1186/s40170-016-0146-8

10 Barile, M., Giancaspero, T. A., Leone, P., Galluccio, M. & Indiveri, C. Riboflavin transport and metabolism in humans. J Inherit Metab Dis 39, 545–557 (2016). 10.1007/s10545-016-9950-0

11 Lienhart, W. D., Gudipati, V. & Macheroux, P. The human flavoproteome. Arch Biochem Biophys 535, 150–162 (2013). 10.1016/j.abb.2013.02.015

12 Martinez-Limon, A., Calloni, G., Ernst, R. & Vabulas, R. M. Flavin dependency undermines proteome stability, lipid metabolism and cellular proliferation during vitamin B2 deficiency. Cell Death Dis 11, 725 (2020). 10.1038/s41419-020-02929-5

13 Nakamura, T. et al. Integrated chemical and genetic screens unveil FSP1 mechanisms of ferroptosis regulation. Nat Struct Mol Biol 30, 1806–1815 (2023). 10.1038/s41594-023-01136-y

14 Kagan, V. E. et al. Oxidized arachidonic and adrenic PEs navigate cells to ferroptosis. Nat Chem Biol 13, 81–90 (2017). 10.1038/nchembio.2238

15 Tan, A. et al. Plasma riboflavin concentration as novel indicator for vitamin-B2 status assessment: suggested cutoffs and its association with vitamin-B6 status in women. P Nutr Soc 79, E658–E658 (2020). 10.1017/S0029665120006072

16 Vande Voorde, J., et al. Improving the metabolic fidelity of cancer models with a physiological cell culture medium. Sci Adv 5, eaau7314 (2019). 10.1126/sciadv.aau7314

17 Cantor, J. R. et al. Physiologic Medium Rewires Cellular Metabolism and Reveals Uric Acid as an Endogenous Inhibitor of UMP Synthase. Cell 169, 258–272 e217 (2017). 10.1016/j.cell.2017.03.023

18 Pedrolli, D. B. et al. The antibiotics roseoflavin and 8-demethyl-8-amino-riboflavin from Streptomyces davawensis are metabolized by human flavokinase and human FAD synthetase. Biochem Pharmacol 82, 1853–1859 (2011). 10.1016/j.bcp.2011.08.029

19 McNulty, H., Pentieva, K. & Ward, M. Causes and Clinical Sequelae of Riboflavin Deficiency. Annu Rev Nutr 43, 101–122 (2023). 10.1146/annurev-nutr-061121-084407

20 Alborzinia, H. et al. LRP8-mediated selenocysteine uptake is a targetable vulnerability in MYCN-amplified neuroblastoma. EMBO Mol Med, e18014 (2023). 10.15252/emmm.202318014

21 Dos Santos, A. F., Fazeli, G., Xavier da Silva, T. N. & Friedmann Angeli, J. P. Ferroptosis: mechanisms and implications for cancer development and therapy response. Trends Cell Biol (2023). 10.1016/j.tcb.2023.04.005

22 Chen, Z. et al. PRDX6 contributes to selenocysteine metabolism and ferroptosis resistance. Mol Cell 84, 4645–4659 e4649 (2024). 10.1016/j.molcel.2024.10.027

23 Hemasa, A., Spry, C., Mack, M. & Saliba, K. J. Mutation of the Plasmodium falciparum Flavokinase Confers Resistance to Roseoflavin and 8-Aminoriboflavin. ACS Infect Dis 10, 2939–2949 (2024). 10.1021/acsinfecdis.4c00289

24 Bjelakovic, G., Nikolova, D. & Gluud, C. Antioxidant supplements to prevent mortality. JAMA 310, 1178–1179 (2013). 10.1001/jama.2013.277028

25 Kontush, A., Finckh, B., Karten, B., Kohlschutter, A. & Beisiegel, U. Antioxidant and prooxidant activity of alpha-tocopherol in human plasma and low density lipoprotein. J Lipid Res 37, 1436–1448 (1996).

26 Rimm, E. B. et al. Vitamin E consumption and the risk of coronary heart disease in men. N Engl J Med 328, 1450–1456 (1993). 10.1056/NEJM199305203282004

## Methods References

1 Dehairs, J., Talebi, A., Cherifi, Y. & Swinnen, J.V. CRISP-ID: decoding CRISPR mediated indels by Sanger sequencing. Sci Rep 6, 28973 (2016).

2 Eggers, L.F. & Schwudke, D. Liquid Extraction: Folch. in Encyclopedia of Lipidomics (ed. Wenk, M.R.) 1–6 (Springer Netherlands, Dordrecht, 2016).

3 Ni, Z., Angelidou, G., Hoffmann, R. & Fedorova, M. LPPtiger software for lipidome-specific prediction and identification of oxidized phospholipids from LC-MS datasets. Scientific Reports 7, 15138 (2017).

4 Criscuolo, A. et al. Analytical and computational workflow for in-depth analysis of oxidized complex lipids in blood plasma. Nat Commun 13, 6547 (2022).

5 Adams, K.J. et al. Skyline for Small Molecules: A Unifying Software Package for Quantitative Metabolomics. J Proteome Res 19, 1447–1458 (2020).

6 Chong, J., Wishart, D.S. & Xia, J. Using MetaboAnalyst 4.0 for Comprehensive and Integrative Metabolomics Data Analysis. Curr Protoc Bioinformatics 68, e86 (2019).

7 Muller, T. et al. Automated sample preparation with SP3 for low-input clinical proteomics. Mol Syst Biol 16, e9111 (2020).

8 Burger, B., Vaudel, M. & Barsnes, H. Importance of Block Randomization When Designing Proteomics Experiments. J Proteome Res 20, 122–128 (2021).

9 Bruderer, R. et al. Extending the limits of quantitative proteome profiling with data-independent acquisition and application to acetaminophen-treated three-dimensional liver microtissues. Mol Cell Proteomics 14, 1400–10 (2015).

10 Pang, Z. et al. MetaboAnalyst 5.0: narrowing the gap between raw spectra and functional insights. Nucleic Acids Res 49, W388–W396 (2021).

11 Zhou, Y. et al. Metascape provides a biologist-oriented resource for the analysis of systems-level datasets. Nat Commun 10, 1523 (2019).

12 Feng, S. et al. The crystal structure of human ferroptosis suppressive protein 1 in complex with flavin adenine dinucleotide and nicotinamide adenine nucleotide. MedComm (2020) 5, e479 (2024).

13 Sastry, G.M., Adzhigirey, M., Day, T., Annabhimoju, R. & Sherman, W. Protein and ligand preparation: parameters, protocols, and influence on virtual screening enrichments. J Comput Aided Mol Des 27, 221–34 (2013).

14 Schrödinger, L. Schrödinger Release 2024-2: Protein Preparation Wizard. (Schrödinger, LLC, New York, NY, 2024).

15 Ryde, U. On the role of Glu-68 in alcohol dehydrogenase. Protein Sci 4, 1124–32 (1995).

16 Case, D.A. et al. AmberTools. J Chem Inf Model 63, 6183–6191 (2023).

17 M. J. Frisch, G.W.T., H. B. Schlegel, G. E. Scuseria, M. A. Robb, J. R. Cheeseman, G. Scalmani, V. Barone, G. A. Petersson, H. Nakatsuji, X. Li, M. Caricato, A. Marenich, J. Bloino, B. G. Janesko, R. Gomperts, B. Mennucci, H. P. Hratchian, J. V. Ortiz, A. F. Izmaylov, J. L. Sonnenberg, D. Williams-Young, F. Ding, F. Lipparini, F. Egidi, J. Goings, B. Peng, A. Petrone, T. Henderson, D. Ranasinghe, V. G. Zakrzewski, J. Gao, N. Rega, G. Zheng, W. Liang, M. Hada, M. Ehara, K. Toyota, R. Fukuda, J. Hasegawa, M. Ishida, T. Nakajima, Y. Honda, O. Kitao, H. Nakai, T. Vreven, K. Throssell, J. A. Montgomery, Jr., J. E. Peralta, F. Ogliaro, M. Bearpark, J. J. Heyd, E. Brothers, K. N. Kudin, V. N. Staroverov, T. Keith, R. Kobayashi, J. Normand, K. Raghavachari, A. Rendell, J. C. Burant, S. S. Iyengar, J. Tomasi, M. Cossi, J. M. Millam, M. Klene, C. Adamo, R. Cammi, J. W. Ochterski, R. L. Martin, K. Morokuma, O. Farkas, J. B. Foresman, and D. J. Fox. Gaussian 09, Revision A.02. (Gaussian, Inc., Wallingford, CT, 2016).

18 Wang, J., Wolf, R.M., Caldwell, J.W., Kollman, P.A. & Case, D.A. Development and testing of a general amber force field. J Comput Chem 25, 1157–74 (2004).

19 Lindorff-Larsen, K. et al. Improved side-chain torsion potentials for the Amber ff99SB protein force field. Proteins 78, 1950–8 (2010).

20 Abraham, M.J. et al. GROMACS: High performance molecular simulations through multi-level parallelism from laptops to supercomputers. SoftwareX 1–2, 19–25 (2015).

21 Jorgensen, W.L., Chandrasekhar, J., Madura, J.D., Impey, R.W. & Klein, M.L. Comparison of simple potential functions for simulating liquid water. The Journal of Chemical Physics 79, 926–935 (1983).

22 Bussi, G., Donadio, D. & Parrinello, M. Canonical sampling through velocity rescaling. J Chem Phys 126, 014101 (2007).

23 Bernetti, M. & Bussi, G. Pressure control using stochastic cell rescaling. J Chem Phys 153, 114107 (2020).

24 Hess, B., Bekker, H., Berendsen, H.J.C. & Fraaije, J.G.E.M. LINCS: A linear constraint solver for molecular simulations. Journal of Computational Chemistry 18, 1463–1472 (1997).

25 Miyamoto, S. & Kollman, P.A. Settle: An analytical version of the SHAKE and RATTLE algorithm for rigid water models. Journal of Computational Chemistry 13, 952–962 (1992).

26 Darden, T., York, D. & Pedersen, L. Particle mesh *Ewald: An N⋅log(N) method for Ewald sums in large systems*. The Journal of Chemical Physics 98, 10089–10092 (1993).

27 Cheatham, T.E., III, Miller, J.L., Fox, T., Darden, T.A. & Kollman, P.A. Molecular Dynamics Simulations on Solvated Biomolecular Systems: The Particle Mesh Ewald Method Leads to Stable Trajectories of DNA, RNA, and Proteins. Journal of the American Chemical Society 117, 4193–4194 (1995).

